# Intracellular Neutralization of Ricin Toxin by Single Domain Antibodies Targeting the Active Site Pocket

**DOI:** 10.1101/805754

**Authors:** Michael J. Rudolph, Timothy F. Czajka, Simon A. Davis, Chi My Thi Nguyen, Xiao-ping Li, Nilgun E. Tumer, David J. Vance, Nicholas J. Mantis

## Abstract

The extreme potency of the plant toxin, ricin, is due to its enzymatic subunit, RTA, which inactivates mammalian ribosomes with near perfect efficiency. Here we characterized, at the functional and structural levels, seven alpaca single-domain antibodies (V_H_Hs) previously reported to recognize epitopes in proximity to RTA’s active site. Three of the V_H_Hs, V2A11, V8E6 and V2G10, were potent inhibitors of RTA *in vitro* and protected Vero cells from ricin when expressed as intracellular antibodies (“intrabodies”). Crystal structure analysis revealed that the complementarity-determining region 3 (CDR3) elements of V2A11 and V8E6 penetrate RTA’s active site and interact with key catalytic residues. V2G10, in contrast, sits atop the enzymatic pocket and occludes substrate accessibility. The other four V_H_Hs also penetrated/occluded RTA’s active site, but lacked sufficient binding affinities to outcompete RTA-ribosome interactions. Intracellular delivery of high-affinity, single-domain antibodies may offer a new avenue in the development of countermeasures against ricin toxin.

## Introduction

Ricin is an extremely potent biological toxin derived from the castor bean (*Ricinus communis*). It has a long history in the development of immunotoxins aimed at combating B cell lymphomas and other cancers (*1*). Yet, ricin is equally renowned as a biothreat agent, especially if dispersed by aerosol (*2*). Ricin’s galactose/N-acetylgalactosamine-binding lectin subunit, RTB, mediates toxin endocytosis and retrograde transport to the endoplasmic reticulum (ER) of mammalian cells. In the ER lumen, ricin’s enzymatic subunit, RTA, is liberated from RTB and retro-translocated into the cytoplasm where it inactivates ribosomes with remarkable efficiency (*3, 4*). Activation of the ribotoxic stress response (RSR) and multiple stress-activated protein kinase (SAPK) pathways ensue, resulting in the triggering of programed cell death pathways. In the context of the lung, ricin triggers acute lung injury characterized by a massive inflammatory response driven by IL-1, IL-6 and members of the tumor necrosis factor-α (TNF-α) superfamily, triggering destruction of the lung epithelium, vascular leak, and edema (*5*).

The catalytic mechanism by which RTA disables mammalian ribosomes was elucidated three decades ago when the X-ray crystal structure of ricin and its enzymatic activities were resolved more or less simultaneously (*6, 7*). Endo and colleagues demonstrated that RTA is an RNA N-glycosidase (EC 3.2.2.22) that depurinates a single adenosine residue within the sarcin-ricin loop (SRL) of the 28S rRNA, an activity measurable in *in vitro* translation assays (*6*). The SRL, one of the longest conserved stretches of rRNA sequence, makes direct interactions with the GTP-binding domains of elongation factors like EF-Tu and is therefore indispensable for peptide elongation. The depurination reaction is confined to RTA’s active site, a large solvent-exposed cleft on one face of the molecule that accommodates the protruding adenine (<u>A</u>) within the conserved G<u>A</u>GA motif of the mammalian SRL. The five critical residues associated with RTA’s enzymatic activity have been defined by site-directed mutagenesis and include Tyr80, Tyr123, Glu177, Arg180, and Trp211 (*8*). Tyr80 and Tyr123 serve to stabilize the adenine base substrate via a π-stacking network. Arg180 is involved in protonation of the adenine leaving group while Glu177 stabilizes the actual cleavage of the N-glycosidic bond. The role of Trp211 in catalysis remains unknown. These catalytic residues, as well as the chemistry of the SRL depurination reaction is conserved among other members of the ribosome-inactivating protein (RIP) superfamily of toxins, including Shiga toxins 1 (Stx1) and 2 (Stx2) from foodborne *Escherichia coli* (*9*).

With the capacity to inactivate >1500 ribosomes per minute (*10*), RTA’s active site is an obvious target to consider when designing therapeutics to arrest the effects of ricin toxin exposure. In fact, early efforts successfully identified substrate analogues (e.g., pteroic acid, guanine-like compounds) with modest RTA inhibitory activity *in vitro* (*9*), while other groups identified molecules capable of trapping RTA’s active site in a closed conformation (*11*). However, issues related to solubility, limited potency and/or biodistribution have severely curtailed the use of those small molecule inhibitors in cell-based assays and animal models of ricin intoxication (*12*). High-throughput, cell-based screens run in parallel as a complementary means of identifying novel ricin inhibitors yielded compounds that targeted host proteins associated with toxin trafficking and SAPK pathways, but not ricin itself (*13, 14*).

In the past decade, camelid-derived, single-domain antibodies, commonly referred to as V_H_Hs or nanobodies, have received enormous attention for their potential as therapeutics against emerging infectious disease and biothreat agents, including botulinum neurotoxin (BoNT), anthrax toxin, and Shiga toxin (*15–18*). V_H_Hs are small (13-16 kDa) immunoglobulin elements amenable to expression in *E. coli* and surface display on bacteriophage M13. V_H_Hs are also highly soluble and thermostable. Of particular relevance to RTA is the reported propensity of V_H_Hs to target active site clefts and enzymatic pockets, as shown for lysozyme, α-amylase and others (*19, 20*). We recently described a collection of 21 V_H_Hs that bind in immediate proximity to or overlapping with RTA’s active site, as demonstrated by epitope mapping studies using hydrogen deuterium exchange (HDX) (*21, 22*). In this report we have characterized seven of those V_H_Hs and demonstrate that three are potent inhibitors of RTA’s enzymatic activity in *in vitro* assay and when expressed as intracellular antibodies (“intrabodies”) within the cytoplasm of target cells. We then solved X-ray crystal structures of each of the V_H_Hs in complex with RTA, which revealed direct interactions with the catalytic residues associated with depurination of the SRL.

## Results

### Identification of V_H_Hs with potent RTA inhibitory activity

We recently identified, through a strategic series of masking and targeted elutions, a collection of 21 V_H_Hs that recognized spatially-distinct epitopes along the rim of RTA’s active site (Figure S1) (*21, 22*). A total of seven were chosen for further examination, ultimately because we were able to successfully solve the crystal structure of each in complex with RTA (Table 1; Figure S1). Two of the V_H_Hs, V2A11 and V6H8, derive from different alpaca libraries but share high degree of CDR3 primary amino acid sequence identity (69%), possibly indicative of a similar mode of interaction with RTA (Figure S1). Three other V_H_Hs, V8E6, V6A7 and V6A6, constitute a clonal family, as evidenced by >80% identity in CDR3 (Figure S1). In addition, V6D4 was one of only two V_H_Hs among the nearly two dozen previously characterized active site-targeting V_H_Hs with detectable toxin-neutralizing activity in a Vero cell assay (Figure S1). The binding affinities of the seven V_H_Hs for RTA were determined by surface plasmon resonance (SPR) and ranged from 0.3 nM to ∼11 nM (Table 1: Figure S2).

**Table 1.** Summary of V_H_H-RTA binding and interface information

| V<sub>H</sub>H | K<sub>d</sub><sup>b</sup> | PDB<sup>c</sup> | H-bonds<sup>a</sup> |  | SB<sup>d</sup> | SC<sup>e</sup> | Buried Surface Area (Å<sup>2</sup>) |  |  |  |
| --- | --- | --- | --- | --- | --- | --- | --- | --- | --- | --- |
|  |  |  | Total | CDR1/2/3 |  |  | Total | CDR1 | CDR2 | CDR3 |
| **V2A11** | 0.31 | 6OBC | 10 | 2/0/7 | 9 | 0.617 | 1953 | 254 | 290 | 1210 |
| **V6H8** | 4.01 | 6OBE | 6 | 1/0/3 | 8 | 0.684 | 2080 | 401 | 355 | 1257 |
| **V8E6** | 0.42 | 6OBG | 19 | 5/3/8 | 5 | 0.701 | 2367 | 452 | 520 | 1188 |
| **V6A7** | 4.2 | 6OBM | 24 | 8/4/10 | 3 | 0.672 | 2416 | 367 | 493 | 1325 |
| **V6A6** | 10.9 | 6OBO | 20 | 5/3/10 | 3 | 0.657 | 2519 | 475 | 494 | 1323 |
| **V2G10** | 0.28 | 6OCA | 10 | 0/3/7 | 10 | 0.665 | 1965 | 405 | 487 | 1194 |
| **V6D4** | 9.51 | 6OCD | 9 | 0/0/6 | 3 | 0.689 | 1746 | 12 | 0 | 1125 |

<sup>a</sup>, hydrogen bonds; <sup>b</sup>, nM; <sup>c</sup>,PBD ID; <sup>d</sup>, salt bridges; <sup>e</sup>, shape complementarity score

The effect of the seven V_H_Hs on RTA’s capacity to inactivate ribosomes was evaluated in a cell-free, luciferase-based *in vitro* translation (IVT) assay, as described in detail in the Materials and Methods. RTA (EC_90_; 0.75 nM) was mixed with V_H_Hs across a dose range and then added to the IVT cocktail. Luciferase production at 90 min served as the proxy for protein synthesis and ribosome activity. Three V_H_Hs (V2A11, V8E6 and V2G10) had high RTA inhibitory activity (>70%), one (V6A7) had intermediate activity (30-70%), and the remaining three V_H_Hs (V6A6, V6H8 and V6D4) had low or undetectable inhibitory activity (Figure 1A). The ability of the seven V_H_Hs to neutralize RTA was dose-dependent (Figure 1B) and correlated directly with binding affinity (K_D_; Tables 1, S1). The relationship between binding affinity and RTA inhibition is exemplified by the three clonally related V_H_Hs, V8E6, V6A7 and V6A6, whose RTA neutralizing activities (high, medium and low, respectively) tracked inversely with their dissociation constants (0.42 nM, 4.2 nM and 10.9 nM, respectively; Table 1).

**Figure 1.**
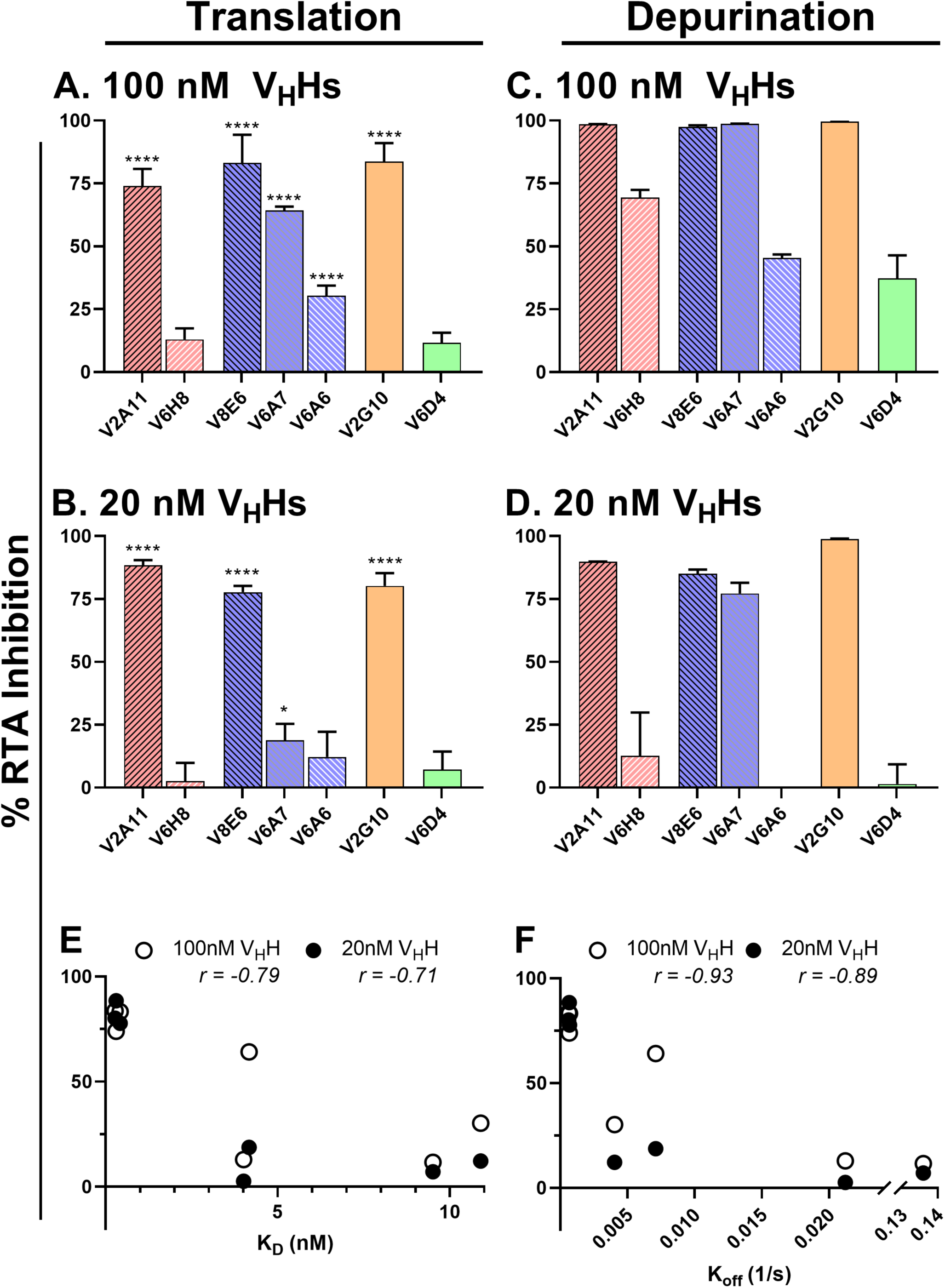
Abrogation of RTA’s RIP activity in vitro by single domain antibodies. **(A, B)** *In vitro* translation assays were performed by mixing RTA (0.75 nM) with **(A)** 100 nM or **(B)** 20 nM of indicated V_H_H and then adding the mixture to a cocktail containing rabbit reticulocyte lysate and luciferase mRNA, as detailed in the Materials and Methods. **(C, D)** Depurination assays were done with purified yeast ribosomes mixed with RTA (1.0 nM) and **(C)** 100 nM or **(D)** 20 nM of indicated V_H_H, as described in the Materials and Methods. For both sets of experiments, results were normalized to RTA treatment without antibody (0%) and reactions without the addition of RTA positive controls (100%). Reactions were performed in triplicate. **(E, F)** Spearman correlation analysis was performed to determine correlation between IVT inhibition and the respective binding affinities **(E)** and off-rates **(F)** for each V_H_H. Significance for the IVT experiments was calculated using a Student’s t-test comparison to the RTA control. **** p<0.0001, *** p<0.001, ** p<0.005, * p<0.05

To confirm that the V_H_Hs actually interfere with RTA’s RNA N-glycosidase activity, we tested the antibodies in an rRNA depurination reaction with yeast ribosomes as substrate (*23*). Overall, the trends were nearly identical to what was observed in the IVT assay: V2A11, V8E6 and V2G10 were potent inhibitors of RTA’s depurination activity, while V6H8, V6A6 and V6D4 were weak inhibitors (Figure 1 C, D). V6A7 and V6H8 inhibited RTA in the depurination assay more effectively than in the IVT assay, possibly reflecting slight differences in the sensitivity and/or stringency of the assays.

In previous studies we and others have reported a relationship between antibody binding affinity and toxin-neutralizing activity, at least in cell-based assays (*22*). Therefore, we performed Spearman correlation analysis to examine the relationship between in vitro RTA inhibition and V_H_H binding affinity at both high and low antibody:RTA molar ratios (Figure 1E,F). There was indeed a correlation between RTA inhibitory activity and V_H_H dissociation constants (K_D_), although a stronger relationship with RTA inhibitory activity and dissociation rates (K_off_). These results indicate that the stability of the antibody-antigen complex is critical for potent inhibition of ricin in cell based killing assays and RTA in *in vitro* assays.

### RTA-specific V_H_H expressed as intrabodies render Vero cells resistant to ricin toxin

With the exception of V6D4, the V_H_Hs described above do not to have toxin-neutralizing activity in a cell-based cytotoxicity assay (Figure S1) (*21, 22*). Nonetheless, we hypothesized that the V_H_Hs, by virtue of the fact that they are postulated to bind epitopes in immediate proximity of RTA’s active site, could protect Vero cells from the effects of ricin if they were successfully expressed in the cell cytoplasm as intracellular antibodies or “intrabodies.” Intrabodies, which refers to the cytosolic or organelle-specific expression of antibody variable regions in the form of single chain Fv (scFv) or V_H_Hs, have been exploited as tools to knock-down intrinsic cytosolic targets like the EGF receptor, as well as extrinsic cytosolic targets like viral antigens and even botulinum neurotoxin (BoNT) (*17, 24*). Theoretically, RTA is an ideal intrabody target, since it is exposed to the cytosolic environment immediately following retro-translocation across the ER membrane.

To test this hypothesis, we constructed pCDNA3.1-based mammalian expression vectors encoding the cDNA for each of the seven RTA-specific V_H_Hs under control of the CMV promoter, essentially as done by Tremblay and colleagues for intracellular expression of BoNT-specific V_H_Hs (*17*). The antibodies were also designed to include a C-terminal E-tag (GAPVPYPDPLEPR) to enable detection in ELISA using commercially available anti-E tag antibodies. The resulting V_H_H-encoding DNA expression vectors were emulsified in liposome nanoparticles (LNP) and used to transiently transfect Vero cells. To assess whether the V_H_Hs were functionally expressed in Vero cells, cell lysates were collected two days following transfection and evaluated by RTA and ricin ELISAs. Expression of six of the seven V_H_Hs was confirmed by ELISA (Figures S3). Only V6D4 was not detected in Vero cell lysates, possibly due to improper folding and/or instability in the intracellular compartment. As will be shown later in this manuscript, V6D4 is the only V_H_H among the seven with a non-canonical disulfide bond between CDR2 and CDR3, which potentially renders the antibody susceptible to the reducing environment of the cytoplasm.

Nonetheless, to examine the capacity of the putative active-site targeting V_H_Hs to protect Vero cells against ricin-induced death, cells were transiently transfected as described above and then challenged with ricin 24 h later (Figure 2A). Vero cell viability was assessed after an additional 48 h incubation period. As controls, Vero cells were transfected with the BoNT-specific V_H_H, ciA-H7 (*17*) or simply mock transfected with empty LNPs. Vero cells were treated with an escalating scale of ricin toxin and a shift in EC_50_ values relative to mock-transfected cells was interpreted as being the result of intracellular V_H_H activity.

**Figure 2.**
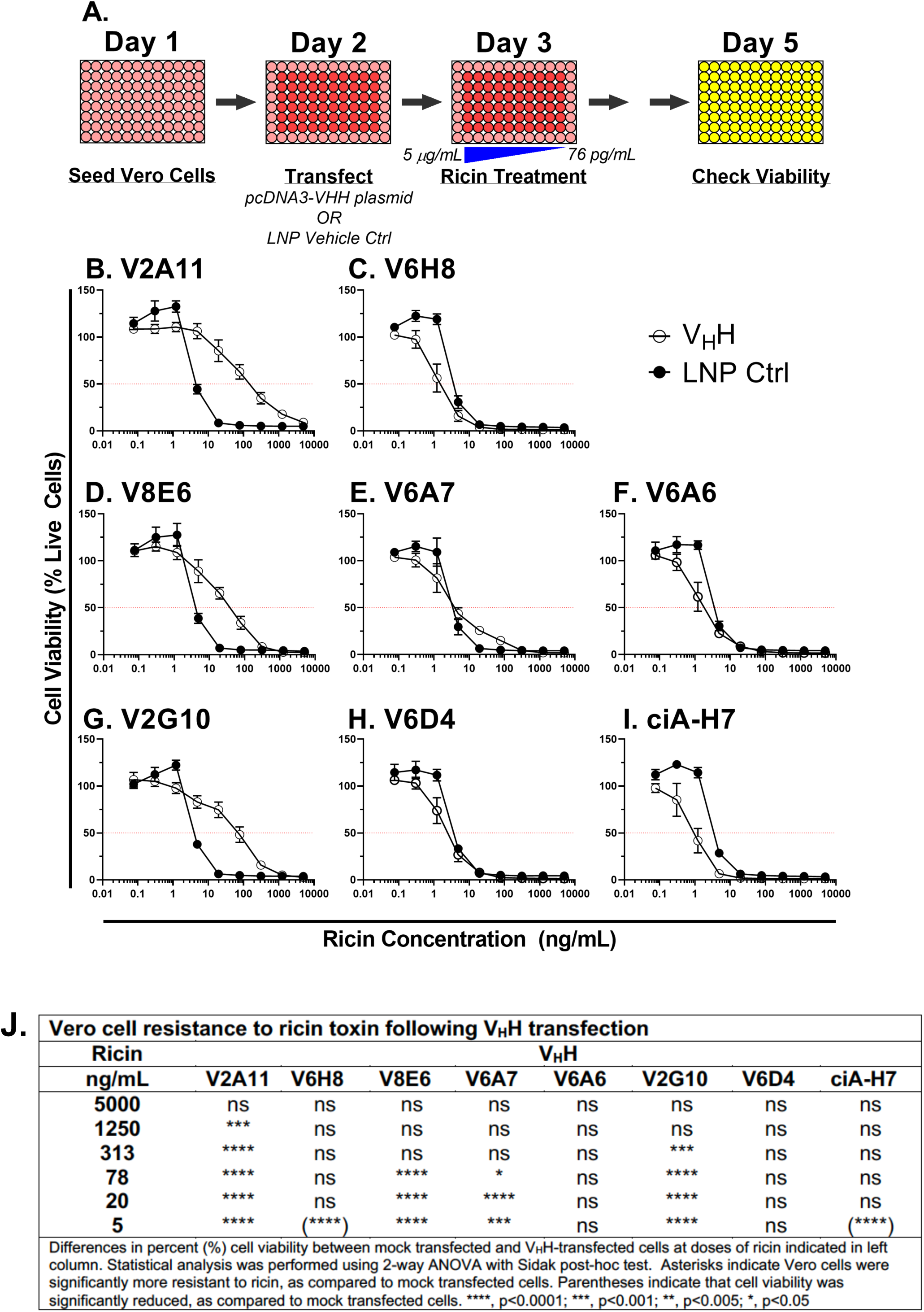
Protection of Vero cells from ricin toxin by V_H_H intrabodies. **(A)** Cartoon of the experimental set-up. Vero cells were seeded in 96-well plates on day 1 and transiently transfected with V_H_H-encoding pcDNA3.1 plasmids on day 2. Ricin challenge occurred on day 3 and cell viability was assessed on day 5. **(B-I)** Cell viabilities of Vero cells transfected with vehicle control (LNP; black circles) or V_H_Hs (open circles) at indicated toxin concentrations. Shown are two biological replicates with three technical replicates each. Horizontal red lines indicate 50% viability. (J) Significance was calculated using a two-way ANOVA with the Sidak correction test at each ricin concentration to compare viability of cells following intrabody transfection and LNP mock transfection. In two instances (indicated by parentheses in panel J), Vero cell viability of V_H_H transfected cells was less than mock transfected cells, suggesting that V_H_H expression actually imparts a burden on cells viability. Asterisks: **** p<0.0001; *** p<0.001; ** p<0.005; * p<0.05.

We found that Vero cells transfected with V_H_Hs V2A11, V8E6, or V2G10 were significantly more resistant to ricin, as compared to mock transfected cells. For example, cells transfected with V2A11 or V2G10 resulted in ∼20-fold increase in EC_50_ values, while transfection with V8E6 resulted in ∼6-fold change (Figure 2B-I). These three V_H_Hs (V2A11, V8E6, or V2G10) were also the most effective at neutralizing RTA *in vitro*, as shown in Figure 1. Transfection with V6A7 rendered the Vero cells resistant to low dose (but not high dose) ricin challenge, whereas the three V_H_Hs with the lowest binding affinities for RTA, namely V6H8, V6A6, and V6D4, afforded no benefit. These results represent the first demonstration of intrabody-mediated targeting of RTA and highlight the potential application of V_H_H-based therapies in combatting other structurally conserved RIPs like Shiga toxins 1 and 2, whose active sites are structurally conserved with RTA (*25*).

### Structural analysis of RTA-V_H_H complexes reveals antibody contacts with active site residues

To better define the molecular basis of intrabody-mediated ricin toxin neutralization, we solved the X-ray crystal structures of each of the seven V_H_Hs in complex with RTA, at resolutions ranging from 1.7 to 2.5 Å (Table 1; Figure 3; Table S2). All seven V_H_Hs possessed the canonical immunoglobulin fold, consisting of nine β-strands arranged in two β-sheets with CDRs 1, 2 and 3 on one face of the molecule along with either 2 or 3 3_10_ helices (*26*). Furthermore, the structure of RTA in each complex consisted of seven α-helices (A-G), three 3_10_ helices (3_10 a_, 3_10 b_, 3_10 c_), and 10 β-strands (a-j) (Figure S4). RTA (PDB: 1RTC) superposition with the seven V_H_H-RTA structures revealed RMSD values ranging from 0.6 to 0.7 Å, demonstrating that RTA does not undergo any significant conformational changes when complexed with the V_H_Hs.

**Figure 3.**
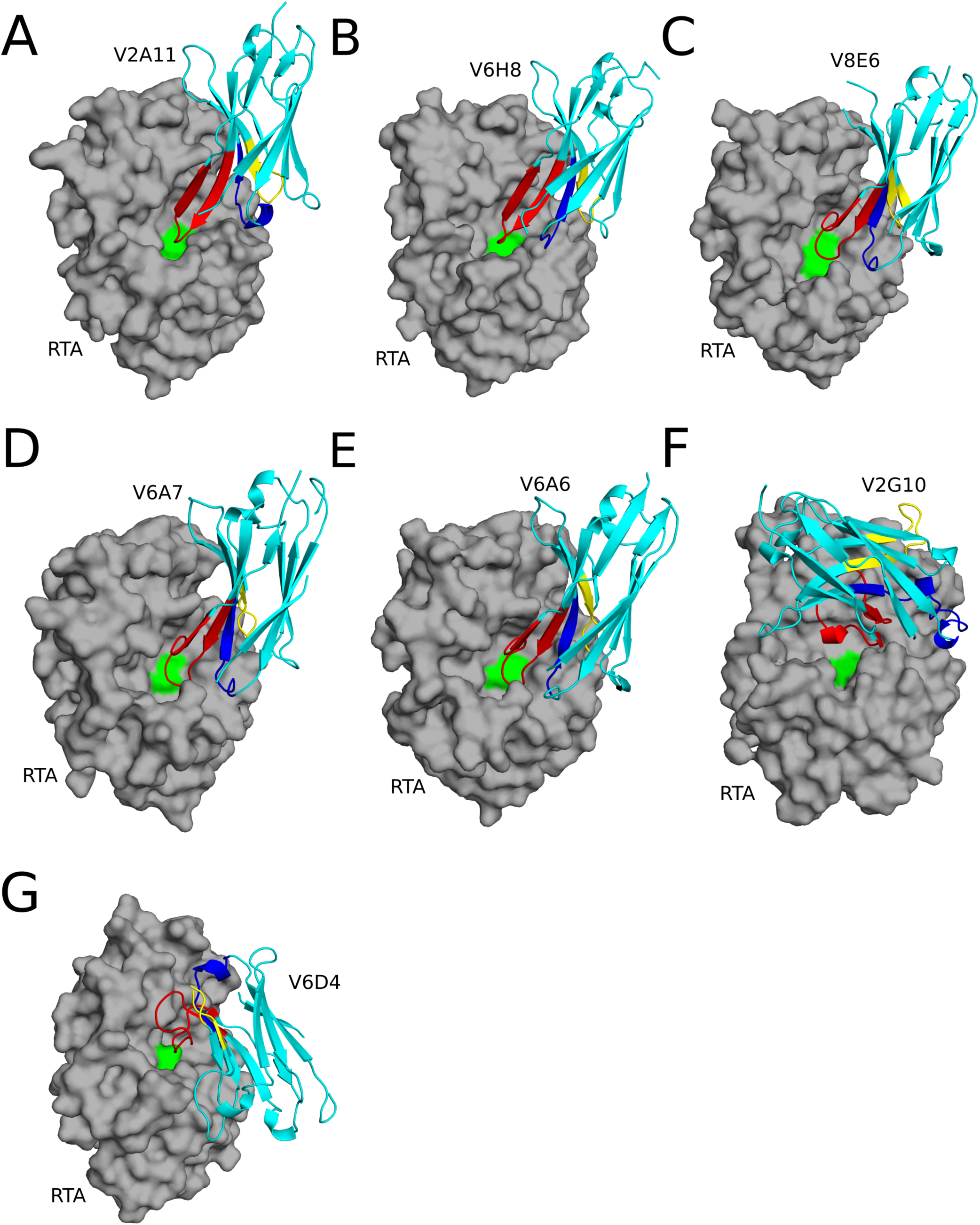
Structures of RTA-V_H_H complexes. Structures of RTA (gray surface) in complex with (A) V2A11, (B) V6H8, (C) V8E6, (D) V6A7, (E) V6A6 (F) V2G10, and (G) V6D4 depicted as ribbon diagrams. Each V_H_H is colored cyan, with CDRs 1, 2, and 3 colored blue, yellow, and red, respectively. RTA active site residue Tyr-80 is colored green.

Upon initial inspection, the most striking feature of the RTA-V_H_H complexes was that five of seven the antibodies, namely V2A11, V6H8, V8E6, V6A7, and V6A6, physically occupied RTA’s active site by virtue of their CDR3 elements (Figure 3A-E). The five RTA-V_H_H complexes were structurally conserved, as evidenced by RMSD values between 0.5 to 2.0 Å when superpositioned (Figure S5). The five V_H_Hs, referred to as Mode 1, are further divisible into two sub-modes (A, B) based on RMSD. Mode 1A, represented by V2A11 and V6H8, have an RMSD of 1.0 Å when superpositioned (Figure S5), while Mode 1B, represented by V6A6, V6A7, and V6E8 has an RMSD of 0.5 to 0.7 Å when superpositioned (Figure S5). The remaining two V_H_Hs, V2G10 (Mode 2) and V6D4 (Mode 3) form lids or caps over RTA’s active site pocket, thereby distinguishing them from Mode 1. As will be detailed below, V2A11 (Mode 1A), V8E6 (Mode 1B) and V2G10 (Mode 2) are each potent inhibitors of RTA even though they attack the active site with slightly different binding modalities.

### Occupancy of RTA’s Active Site by V2A11 and V6H8 (Mode 1A)

Mode 1A consists of two clonally unrelated V_H_Hs with similar CDR3 sequences: V2A11, a potent inhibitor of RTA, and V6H8, a poor inhibitor of RTA. The V2A11-RTA complex resulted in a buried surface area (bsa) of 1953 Å^2^ and an overall shape complementarity score as measured by the shape correlation (Sc) parameter of 0.617 (Table 1). The interaction is dominated by CDR3, which not only penetrates into RTA’s active site but also interacts with residues integral to ricin’s RNA N-glycosidase activity, Tyr80 and Tyr123, which are involved in stacking the adenine substrate within the SRL. Specifically, V2A11 residue Ser103 sits above RTA Tyr80, while V2A11 residue Trp100 comes within 3.5 Å of RTA Tyr123 (Figures 4A). As a result, V2A11 confines Tyr80 in a conformation that would prevent stacking of an adenine base between itself and Tyr123 (Figure S6). V2A11’s CDR1 and CDR2 have only moderate interactions with RTA, as reflected by bsa of 254 Å^2^ and 290 Å^2^, respectively.

**Figure 4.**
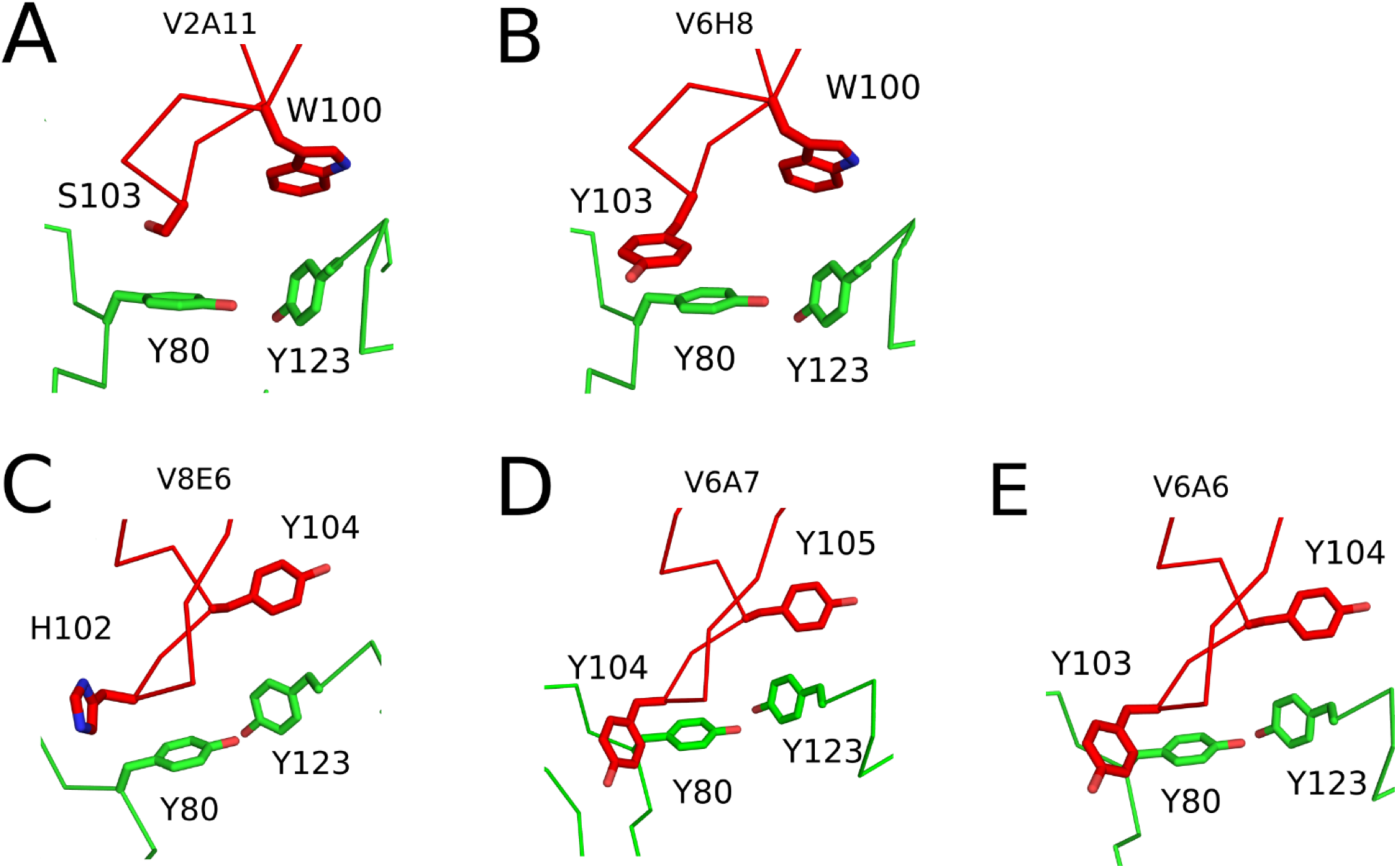
Mode 1 V_H_H interactions within the RTA’s active site residues. The Cα-traces of RTA (green) in complex with (A) V2A11, (B) V6H8, (C) V8E6 (D) V6A7, and (E) V6A6. The V_H_H CDR3 elements are colored red. V_H_H residues forming interactions with RTA active site residues Tyr-80 and Tyr-123 are drawn as sticks. All stick representations are color coordinated to their respective main chain color.

The V6H8-RTA interaction is similar. It results in a total bsa of 2080 Å^2^ and has an Sc score of 0.684 (Table 1). The antibody-toxin interaction is also dominated by a CDR3 that penetrates RTA’s active site. Moreover, V6H8’s Tyr103 obstructs RTA’s Tyr80, while V6H8’s Trp100 comes within 3.9 Å of RTA’s Tyr123 (Figures 4B). CDR1 and CDR2 interactions with RTA results in bsa of 401 Å^2^ and 355 Å^2^, respectively. The discrepancy in RTA binding affinity between V6H8 and V2A11 is conceivably due to a lower entropic penalty endured by V2A11 upon RTA binding relative to V6H8. V2A11’s CDR1 and CDR2 interact to a lesser degree with RTA than V6H8’s CDR1 and CDR2, essentially maintaining greater entropy after RTA interaction. The differences in CDR1 conformation can be explained by several non-covalent interactions unique to V2A11, including a salt bridge between Glu-26 and Lys-99, as well as two hydrogen bonds between the side chain hydroxyl groups of Ser-27 and Ser-29, along with Ser-30’s side chain hydroxyl group with the carbonyl oxygen of Ser-27 (Figure S7A).

Collectively, these non-covalent interactions support the folding of V2A11’s CDR1 residues 27 to 30 into a 3_10_ helix that projects away from RTA, relative to V6H8’s CDR1, ostensibly generating higher configurational entropy within V2A11 when bound to RTA favoring tighter binding. Incidentally, if V2A11’s CDR1 adopted a similar CDR1 configuration as seen in V6H8 when bound to RTA not only would V2A11 lose the aforementioned configurational entropy but V2A11 would also squander two hydrogen bonds formed by Ser-27 with Ser-29 and Ser-30 along with the Glu-26 to Lys-99 salt bridge while forming only one hydrogen bond between Ser-30 and RTA’s Asn-122 as seen with V6H8’s CDR1. This would reduce the enthalpic gain upon RTA binding and also diminish binding affinity for RTA. Moreover, there is a salt bridge between Glu-26 and Lys-99 in V2A11’s CDR1 that cannot form in V6H8 because of a glycine residue at position 26 (Figure S7). The influence of this salt bridge on CDR1 structure is underscored the fact that V6H8’s CDR1 shares structural similarity the CDR1 elements of the three V_H_Hs in mode 1B (V8E6, V6A7, V6A6), which also have a Gly at reside 26 (Figure S7). Finally, V6H8 CDR2 residues 53 to 55 contact RTA to a greater extent than the corresponding residues in V2A11 due to the presence of two salt bridges (CDR2 Arg-55 and RTA Glu-127, and CDR2 Arg-56 and RTA’s Glu-135), which ultimately constrains V2A11 CDR2 and further reduces V2A11’s entropic penalty when binding RTA (Figure S7). Collectively, V2A11’s relative higher binding affinity for RTA, compared to V6H8 can be attributed to a lower overall energetic penalty incurred upon engaging with RTA.

### Occupancy of RTA’s Active Site by V8E6, V6A7 and V6A6 (Mode 1B)

Within Mode 1B, there were notable parallels in how V8E6, V6A7 and V6A6 engaged RTA, which was not surprising considering the three V_H_Hs share 85 to 91% amino acid sequence identity (Figure S1). V8E6 had the strongest binding affinity (K_D_ = 0.42 nM) for RTA and was the most effective at inhibiting RTA *in vitro* and when expressed as an intrabody (Table 1; Figure 2). The co-crystal structure revealed that V8E6 buried 2367 Å^2^ of solvent exposed surface and had the highest shape complementarity in this subgroup with a score of 0.701, accounting for its high binding affinity and potent RTA inhibitory activity (Figure 3C).

V8E6’s CDR3 formed 8 hydrogen bonds with RTA and contributed 1,188 Å 2 of buried surface area (Table 1). Like the two V_H_Hs in described for Mode 1A, V8E6’s CDR3 also occupies RTA’s active site pocket. CDR3 residue Tyr103 obstructs RTA’s Tyr80, while residue Tyr104 obstructs RTA’s Tyr123 (Figure 4C). CDR1 and CDR2 contributions are also significant with 8 H-bonds formed with RTA and a total of 972 Å^2^ of buried surface area.

The co-crystal structure of V6A7-RTA revealed a buried surface area of 2,416 Å^2^ (Table 1;Figure 3D) CDR3 forms 10 hydrogen bonds with RTA and accounts for more than half the total buried surface area (1325 Å^2^). V6A7’s CDR3 also occludes key RTA active site residues: Tyr103 obstructs RTA Tyr80, while Tyr105 comes within 3.8 Å of RTA’s Tyr123 (Figure 4D). As noted in Table 1, V6A7 has a slightly lower shape complementarity with RTA (0.672), as compared to V8E6 (0.701), likely contributing to their different binding affinities (4.2 vs. 0.42 269 nM).

Finally, V6A6 buried the largest amount of solvent exposed surface with RTA (bsa = 2,519 Å^2^), but had the lowest shape complementarity (Sc = 0.657) and the lowest overall binding affinity (10.9 nM) (Table 1). V6A6 established four fewer hydrogen bonds with RTA than V6A7, and two fewer salt bridges with RTA than V8E6, possibly explaining its weak binding affinity for RTA. Nonetheless, V6A6’s CDR3 residues Tyr-103 and Tyr-104 engage active site residues: Tyr-103 occludes RTA residue Tyr80, while Tyr-104 stacks with RTA Tyr123 at a distance of 3.9 Å (Figure 4E). Therefore, binding affinity, rather than epitope specificity, accounts for the failure of V6A6 to inhibit RTA activity in vitro and intracellularly as an intrabody.

### Conserved interactions of Mode 1 V_H_Hs with RTA active site

Taken as a whole, it is remarkable that all five V_H_Hs in Mode 1 employ CDR3 in a similar fashion to occupy RTA’s active site, despite the very limited primary amino acid sequence identity (8-28%) among the CDR3 elements themselves. For example, in all five cases, CDR3 residues 102, 103, or 104 make close contact (2.9 to 3.0 Å) with Tyr80. A second CDR3 residue, Trp100 in V2A11 and V6H8, Tyr104 in V8E6 and V6A6, and Tyr105 in V6A7, are within 3.5 to 3.9 Å of Tyr123. V_H_Hs in Mode 1 also invariably form an H-bond between CDR3 and RTA’s Arg180, the residue responsible for protonating the N-3 atom of the adenine substrate and thereby facilitating hydrolysis of the bond between adenine N-9 and C-1’ of the ribose (*27*).

Ironically, the CDR3 elements also account for the relatively large RMSD (1.4 to 2.0 Å) between Modes 1A and 1B, compared to the RMSD (0.5 to 0.9 Å) within the individual Modes. Specifically, the difference in CDR3 configurations between V_H_Hs in Mode 1A and Mode 1B is determined in large part by the residue located at position 100 (101 in V6A7). In Mode 1A that residue is a Trp, while in Mode 1B that residue is a Gly (Figure S1). A Gly at position 100 is consequential because it permits bulky residues to be positioned at residue 104 or 105 (e.g., Tyr-105 in V6A7; Tyr-104 in V8E6/V6A6), in addition to influencing the spatial location of abutting residues (Thr-102, Tyr-103, and Tyr-104 in V6A7; Ser-101, Tyr-102, and Tyr-103 in V6A6; Ser-101, His-102, and Tyr-103 in V8E6), relative to the corresponding residues in CDR3 of V2A11 and V6H8 (Figure S8B). The CDR3 configurations of Modes 1A and 1B V_H_Hs are also influenced by CDR1 in that in Mode 1A, residue 32 is a Tyr (V2A11) or a His (V6H8), which would sterically clash with a bulky residue located in CDR3 at position 104 or 105 (Figure S5). Altogether, local structural differences in CDR3 configurations result in appreciably different interactions with RTA between V_H_Hs in Modes 1A and 1B.

The CDR1 and CDR2 elements also contributed to the interface with RTA, but strictly outside of RTA’s active site pocket. In general, CDR1 made contact with the loop region between β-strand h and α-helix C (referred to as loop h-C) while CDR2 interacted primarily with α-helix C, the loop between α-helices C-D, and helix 3_10 b_. Altogether, the level of interaction between each CDR1 and CDR2 differed to some degree with V2A11’s CDR1 and CDR2 contributing the least (bsa = 544 Å^2^, 2 H-bonds) and V8E6’s CDR1 and CDR2 contributing the most (bsa = 972 Å^2^, 8 H-bonds) to the interface with RTA. Additional contacts with RTA made by each CDR3 in Mode 1 involved comparable interactions with the α-helices B, C, E and G, β-strands f, i and j, helix 3_10 b_, as well as the loop regions between β-strand g and α-helix B (referred to as loop g-B), α-helices C and D (referred to as loop C-D), β-strands b-c (referred to as loop b-c) and β-strands e-f (referred to as loop e-f) (Table 1).

### Interactions of V2G10 and V6D4 with RTA’s active site

We next analyzed the co-crystal structures of the remaining two complexes, V2G10-RTA and V6D4-RTA. V2G10, a potent inhibitor of RTA, and V6D4, a weak inhibitor of RTA, each formed a cap over RTA’s active site (Figure 3F-G). Compared to the five V_H_Hs in Mode 1, V2G10 and V6D4 are rotated counterclockwise by ∼99° and ∼165°, respectively, around RTA’s active site (Figure S8). In addition, their CDR3 elements are tethered to the surface of the V_H_H molecule, physically restricting their ability to project into the active site cleft. In V2G10, tethering is the result of hydrophobic interactions between CDR3 residues Ala-118 and Tyr-124 and framework region (FR) residues Phe-44 and Phe-55, as well as a salt bridge between CDR3 Arg-119 and FR residue Asp-71. In the case of V6D4, the CDR3 element is restrained by a disulfide bond between CDR2 Cys50 and CDR3 Cys105 (Figure S8).

Overall, the V2G10-RTA interaction buried 1,965 Å^2^ with an Sc score of 0.665 (Table 1). V2G10’s CDR1 and CDR2 elements established 3 hydrogen bonds with RTA and bury a total of 892 Å^2^ of surface area. CDR1 makes contacts with RTA α-helix C, helix 3_10 b_, and β-strands i and j, while CDR2 interacts primarily with α-helix G and β-strands i and j. CDR3 contacts RTA’s α-helices B, C, and G, helix 3_10 b_, β-strand i, and the g-B loop, buries 1194 Å^2^ of solvent exposed surface area, and forms a total of 7 hydrogen bonds with RTA. The CDR3 comes within 12 Å^2^ of the catalytic residue, Tyr-80 (Figure 5A).

**Figure 5.**
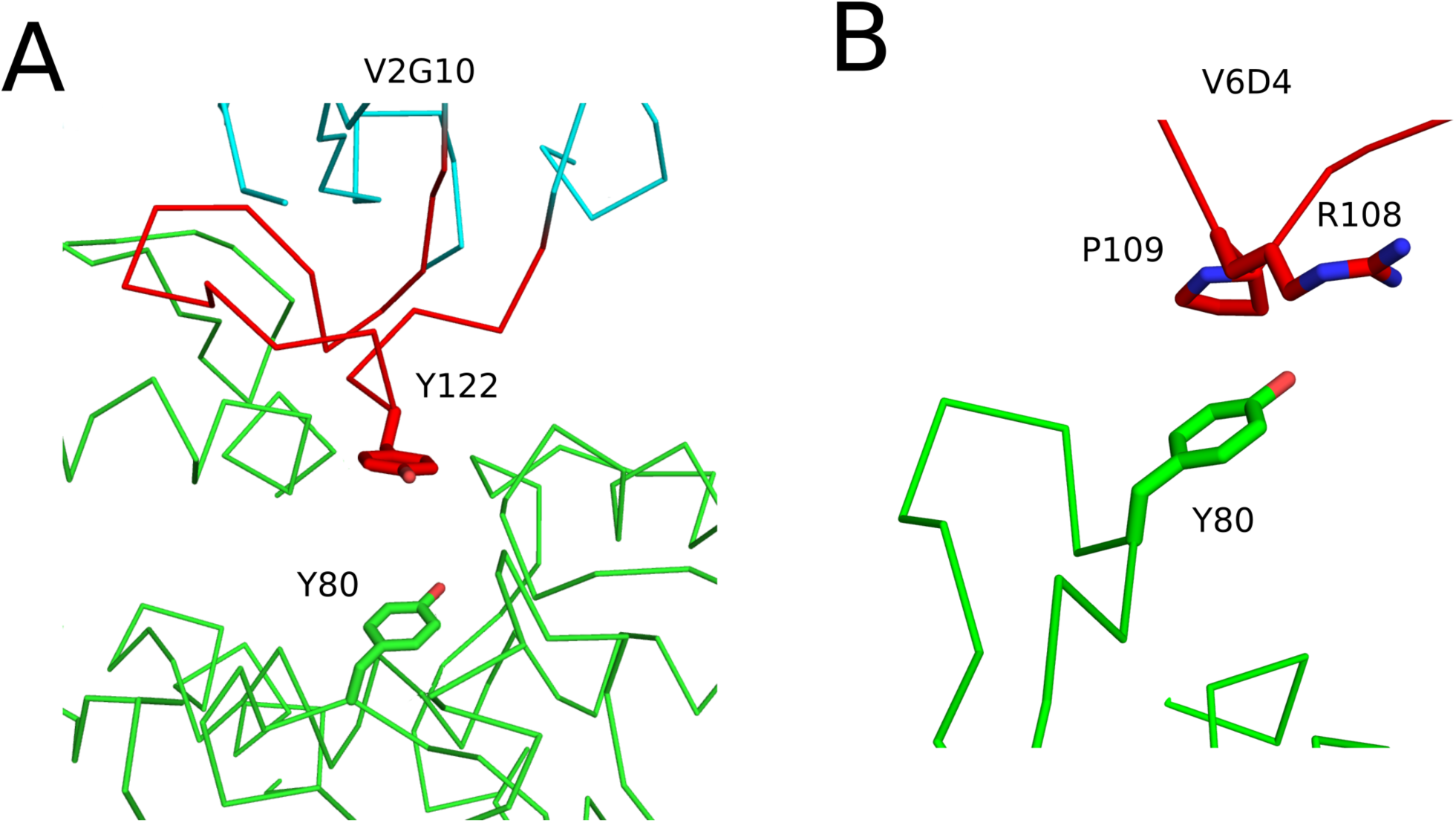
Structural analysis of V6D4 and V2G10 with RTA’s active site residues. Close-up images of the interface between RTA (green) and (A) V2G10 and (B) V6D4. The V_H_Hs CDR3 elements are colored red. V_H_H residues forming interactions with RTA active site residues Tyr-80 are drawn as sticks. All stick representations are color coordinated to their respective main chain color.

The V6D4-RTA interaction generated six hydrogen bonds and buried 1,746 Å^2^ of surface area with an overall Sc of 0.689 (Table 1). V6D4’s CDR1 and CDR2 made minimal contact with RTA, burying 12 Å^2^ and no hydrogen bonds. In contrast, CDR3 buried 1,125 Å^2^ and formed 6 hydrogen bonds. CDR3 interacted with RTA α-helices B, C, and G as well as loops c-d and e-f. In an interesting twist, the presence of a proline residue (Pro-106) in the proximal region of CDR3 creates a partially unrestricted bend within the V_H_H main chain that positions Arg108 and Pro109 within ∼3.5 Å of Tyr80 (Figure 5B).

The crystal structure of V6D4-RTA revealed that V6D4 FR residues (Tyr-45, Gly-46, Tyr-59) and one CDR3 residue (Ile-107) made contact with RTA’s α-helix B (Asn-97, Gln-98, Ala-101) burying 324 Å^2^ while forming 1 hydrogen bond between FR residue Tyr-59 and Asn-97 in RTA’s α-helix B (Figure S9). Although V6D4 is a weak inhibitor of RTA *in vitro*, it was the only V_H_H among the seven that had demonstrable toxin-neutralizing activity in a cell-based assay (Figure S1), which we speculate may be due to contact with RTA’s α-helix B, a known neutralizing hotspot. In this regard, it is interesting to point out that previous epitope mapping studies by HX-MS did not differentiate V2G10 from the Mode 1 family of V_H_Hs, but did successfully classify V6D4 as being unique, largely because of its interaction with α-helix B (*21*).

### Evidence for distinct germline-dictated CDR3 configurations within alpaca derived V_H_Hs

A closer examination of CDR3 configurations between Mode 1 (V2A11, V6H8, V6A6, V6A7, V8E6) and Modes 2 (V2G10) and 3 (V6D4) is warranted, because these groups of V_H_Hs appear to be representative of two CDR3 archetypes that exist in alpacas (*Vicugna pacos*). Mode 1 antibodies like V2A11 are representative of a class of V_H_Hs in which the CDR3 element projects away from the antibody surface in an “extended” conformation, while V2G10 is representative of V_H_Hs in which the CDR3 is “pinned” to the surface of the antibody (Table S3). While our sample size remains small (seven deposited to PDB, plus six pending for “extended”; 11 deposited to PDB, plus 10 pending for “pinned”), the overall alignment is remarkable (Table S3; Figure S10). Within our collection, the pinned-back CDR3 elements are, on average, 5 residues longer than the extended CDR3 elements. Moreover, the two different CDR3 configurations appear to be influenced by germline V_H_ gene usage: the extended elements predominantly derive from IGHV3S53 and the pinned-back from IGHV3-3 (*28*). The fact that V2A11, V8E6 and V2G10 are equally effective at occluding RTA’s active site using divergent molecular approaches to RTA’s active site is a demonstration of the versatility of antibody selection.

## DISCUSSION

RTA is a near perfect enzyme that once unleashed in the cytoplasm of mammalian cells inactivates ribosomes with an efficiency theoretically limited only by diffusion (*3, 6, 12*). RTA’s active site is a solvent exposed cleft on one side of the molecule that accommodates the protruding adenine (<u>A</u>) within the conserved G<u>A</u>GA motif of the mammalian SRL. The adenine base is stabilized within RTA’s active site by a π-stacking network involving Tyr80 and Tyr123. RTA residues Glu177 and Arg180 promote the actual cleavage of the N-glycosidic bond that liberates the adenine moiety, thereby inactivating the ribosome. While RTA’s active site is an attractive drug target, in silico-based efforts to identify small molecule inhibitors of ricin have been stymied on several fronts (*12*). Even the best substrate analog inhibitors to date have IC_50_s only in the micromolar range in cell-free assays. Moreover, while pterin-based molecules that fit snugly into RTA’s active site have been identified, the compounds themselves are poorly soluble and incompatible with cell and animal models of ricin toxicity (*12, 29*). In an effort to circumvent the issue of solubility, high-throughput, cell-based screens were performed with commercially available small molecule libraries. While compounds were identified that were capable of protecting cells from ricin-induced killing, the molecules invariably targeted host cell processes, rather than the toxin itself (*13, 14, 30*).

It is therefore highly significant that we have discovered three V_H_Hs with potent RTA inhibitory activity: V2A11, V8E6 and V2G10. Two of the three antibodies, V2A11 and V8E6, representing binding Modes 1A and 1B, respectively, physically occupy RTA’s active site by virtue of their CDR3 elements, as revealed by the X-ray crystal structure of the V_H_H-RTA complex. In this respect, V2A11 and V8E6 are textbook examples of the propensity of V_H_Hs to target concave epitopes such as the enzymatic pockets of hen egg white lysozyme and α-amylase (*19, 20, 31-33*), the CD4-binding pocket of HIV-1 gp120 (*34*), and the receptor-binding pocket of poliovirus (*35*). The proclivity of camelid heavy chain-only antibodies for enzymatic clefts has been attributed to their “prolate” shape and rigidification of CDR3 through FR interactions, as well as non-canonical disulfide bonds that alleviate the entropic penalty that would normally be associated with extended CDR3 elements (*15, 33*). The interactions of V_H_Hs with active site pockets can be so exquisite that they compete with small molecule substrate analogs and mimic native substrates at the atomic level (*19, 20, 36*).

Indeed, we found that V_H_Hs in Mode 1A and 1B had CDR3 residues that directly interacted with RTA’s catalytic residues Tyr80 and Tyr123, steering Tyr80 into a configuration that would effectively perturb the π-stacking network required for depurination of SRL (Figure S6). Mode 1 V_H_H CDR3 elements further perturbed RTA’s RNA N-glycosidase machinery by forming H-bonds with Arg180, effectively pulling this residue away from the target N-3 atom within the adenine ring. By way of comparison, when RTA is bound to a substrate analog, AMP, Arg-180 is 3Å from the N-3 atom (PDB: 3HIO), whereas Arg-180 is 3.9 Å away from the N-3 atom when RTA is bound to V2A11 (Figure S6). The greater distance between Arg-180 and substrate when RTA is bound to Mode 1 V_H_Hs would compromise proton transfer to the N-3 atom within the adenine ring, effectively impeding hydrolysis of the N-9 to C-1’ bond within the adenosine residue within the SRL substrate (*27*).

The other three V_H_Hs within Mode 1, V6H8, V6A7 and V6A6, engaged with RTA active site residues in an identical fashion as V2A11 (in the case of V6H8) and V8E6 (in the case of V6A7 and V6A6), but were ineffective as RTA inhibitors. The primary difference between the potent and impotent antibodies is binding kinetics: V2A11 and V8E6 have equilibrium dissociation constants (K_D_) of <1 nM and off-rates of <6 x 10^-4^ (1/s), whereas V6H8, V6A7 and V6A6 have K_D_ values of 4-10 nM and off-rates ranging from 4-20 x 10^-3^ (1/s). These results reveal a minimum binding/dissociation threshold necessary to effectively neutralize RTA, a topic that will be discussed further below.

The third potent inhibitor of ricin, namely V2G10, sits atop and caps (rather than penetrates) RTA’s active site. This alternative mode of attack (vis a vis V2A11 and V8E6) is due to V2G10’s CDR3 element being tethered to the surface of the antibody and, therefore, restricted from projecting away from the core immunoglobulin fold and interacting with concave epitopes (*33*). We speculate that the pinned back CDR3 configuration is predetermined by the IGHV3-3 VH gene usage and represents one of at least two alpaca V_H_H archetypes. Despite these different binding modalities, V2G10’s K_D_ (0.28 nM) and off-rate (6.2 x 10^-4^ 1/s) were identical to those of V2A11 and V8E6 and therefore over the minimum threshold required to inhibit RTA activity.

All three potent inhibitors of RTA, V2G10, V2A11 and V8E6, also rendered Vero cells resistant to ricin intoxication when expressed as cytoplasmic “intrabodies”. While intrabodies in the form of single chain Fv (scFv) or V_H_Hs have been under investigation for more than a decade as tools to deplete (“knockdown”) viral and oncogenic proteins in mammalian cells, there is only a handful of examples targeting toxins, and only one instance aimed at a toxin’s enzymatic activity (*37*). Specifically, Tremblay and colleagues identified two V_H_Hs against botulinum neurotoxin (BoNT) serotype A that blocked the toxin’s protease activity *in vitro* (*17*). When expressed as intrabodies in neuronal cells, the V_H_Hs co-localized with the toxin and reduced protease-mediated degradation of BoNT/A’s substrate, SNAP25. At this point in time, we speculate that V2G10, V2A11 and V8E6 exert their effects on ricin toxin by interacting with RTA immediately following its translocation across the ER membrane and preventing subsequent interactions with the ribosomes. The window of opportunity for the antibodies to intervene may be extremely short, considering that there is evidence that ribosomes promote RTA refolding and stabilization (*38*). Moreover, the results presented here certainly argue that high affinity binding (slow off-rates) are pre-requisite for intracellular anti-toxin activity. With the advent of mRNA-based, V_H_H-expression platforms for passive vaccination (at least in mice) (*39*), the prospect of using V_H_Hs as therapeutics for fast acting biothreat agents is no longer farfetched.

## Experimental Design

### Cloning, expression and purification of V_H_Hs

The PCR amplicons for all 7 V_H_Hs were subcloned into the pSUMO expression vector which contained an N-terminal deca-histidine and SUMO tag, as described (*40*). RTA (residues 1-268) was subcloned into the N-terminally deca-histidine tagged MCSG7 expression vector. All cloning was performed using a standard ligase independent cloning protocol. The V_H_Hs and RTA were expressed in *E. coli* strain BL21(DE3)-pRARE. The transformed bacteria were grown at 37°C in TB medium and induced with IPTG (0.1 mM) at 20°C and grown for ∼16 h to mid-log phase (optical density of 0.6 at 600 nm). Cells were harvested by centrifugation and resuspended in 20 mM HEPES [pH 7.5] and 150 mM NaCl. The cell suspension was sonicated and centrifuged at 30,000 *g* for 30 min. The supernatants were subjected to 1 mL nickel affinity column followed by a Superdex 200 16/60 gel filtration column on an AKTAxpress (GE Healthcare). The elution buffer consisted of 0.5M imidazole in binding buffer. The gel filtration buffer consisted of 20 mM HEPES [pH 7.5], 150 mM NaCl, and 20 mM imidazole. V_H_H- or RTA-containing fractions were pooled and subject to TEV protease cleavage (1:10 weight ratio) for 3 h at room temperature in order to remove fusion protein tags. The cleaved protein was passed over a 1 mL Ni-NTA agarose (Qiagen) gravity column to remove TEV protease, cleaved residues, and uncleaved fusion protein. To generate each RTA-V_H_H protein complexes, RTA was mixed with V_H_Hs in a 1:1 stoichiometry then concentrated to of 10 mg/ml.

### *In vitro* measures of RTA enzymatic activity

IVT assays were performed as described, with minor alterations (*41*). V_H_Hs diluted into 1% bovine serum albumin in PBS (w/v) were mixed at defined stoichiometric ratios with 0.75 nM RTA (Vector Laboratories, Burlingame, CA) in triplicate then added to an ice cold mixture containing luciferase mRNA (3.7 μg/mL; Ambion, Inc., Austin, TX) and Retic Lysate IVT™ Kit (ThermoFisher Scientific). The cocktail was incubated for 90 min at 30°C and then chilled on ice for 5 min before being transferred to wells of an opaque, 96-well microtiter plate with an equal volume (20 μL) of Bright-Glo™ luciferase substrate (Promega, Madison, WI) at room temperature. Luminescence was detected immediately using SpectraMax L (Molecular Devices, San Jose, CA). Percentage of RTA inhibition was scaled with 0 at the average luminescence of RTA control reactions and 100 at the average luminescence of positive control reactions.

The depurination inhibition activities of V_H_Hs were measured by qRT-PCR (*42*). Briefly RTA (1.0 nM) was mixed with 4, 20 or 100 nM V_H_Hs in 100 µL volume at room temperature. Yeast ribosomes (50 nM) were added and the mixture was incubated at 25 °C for 5 min. The reaction was stopped by the addition of 100 µL 2X RNA extraction buffer. RNA was purified from the reaction mixture and the depurination was measured using a previously published qRT-PCR method (*43*). For each set of reactions, one sample without the addition of RTA was set as the 100% inhibition control and one sample without the addition of V_H_H was set as the no inhibition control. The percentage of depurination inhibition was calculated based on the controls. The data is shown as mean ± SD from three different biological replicates.

### Crystallization and data collection

All RTA-V_H_H crystals were grown by sitting drop vapor diffusion at 20°C using a protein to reservoir volume ratio of 1:1 with total drop volumes of 0.4 μL. Crystals of the RTA-V2A11 complex were produced using a crystallization solution containing 100 mM Tris (pH 7.0), 200 mM calcium acetate, and 20% PEG 3,000. Crystals of the RTA-V2G10 complex were produced using a crystallization solution containing 100 mM MES (pH 5.0) and 20% PEG 6,000. Crystals of the RTA-V1D3 complex were produced using a crystallization solution containing 100 mM sodium acetate (pH 4.6) and 1.1 M di-ammonium tartrate. Crystals of the RTA-V8E6 complex were produced using a crystallization solution containing 100 mM MES (pH 6.0), 170 mM zinc acetate, 6% PEG 8,000, and 0.5% ethyl acetate. Crystals of the RTA-V6A7 complex were produced using a crystallization solution containing 200 mM potassium sulfate and 20% PEG 3,350. Crystals of the RTA-V6H8 complex were produced using a crystallization solution containing 100 mM citric acid and 15% PEG 6,000. Crystals of the RTA-V6D4 complex were produced using a crystallization solution containing 100 mM MES (pH 6.0), 400 mM zinc acetate, and 11% PEG 8,000. Crystals of the RTA-V6A6 complex were produced using crystallization buffer containing 100 mM imidazole (pH 8.0), 200 mM calcium acetate, and 15% PEG 8,000. All crystals were flash frozen in liquid nitrogen after a short soak in the appropriate crystallization buffers supplemented with 25% ethylene glycol. Data were collected at the 24-ID-C and 24-ID-E beamlines at the Advanced Photon Source, Argonne National Labs as well as beamline 8.2.2 at the Advanced Light Source, Berkeley Lab. All data was indexed, merged, and scaled using HKL2000 (*44*) then converted to structure factors using CCP4 (*45*).

### Structure determination and refinement

The RTA-VHH complex structures were solved by molecular replacement using the program Phaser (*46*). Molecular replacement calculations were performed using the coordinates of the ricin a chain as a search model for RTA (PDB code 1RTC) and the VHH coordinates D10 as the search model for each VHH (PDB code 3EZJ). The resulting phase information from molecular replacement was used to autobuild the polypeptide chain for each RTA-VHH complex using the program ARP (*47*). Further manual model building was performed with COOT (*48*)combined with structural refinement employing the PHENIX package (*49*). Twinned refinement was performed for the RTA-V2G10 complex using the twin operator -h-2*l,-k,l with a twinning fraction of 0.052. During refinement a cross-validation test set was created from a random 5% of the reflections. Data collection and refinement statistics are listed in Table S1. Molecular graphics were prepared using PyMOL(Schrodinger) (DeLano Scientific LLC, Palo Alto, CA).

### Accession Numbers

The structures generated in this study were deposited in the Protein Data Bank (PDB; http://www.rcsb.org/pdb/). The accession numbers are presented in Table 1 and Table S1.

### Surface Plasmon Resonance (SPR)

Rabbit anti-E-tag polyclonal antibody (Bethyl Laboratories #A190-133A) was immobilized at 9000 RU on a Series S CM5 Chip. V_H_Hs with a C-terminal E-tag (diluted in running buffer) were captured on the chip surface at a maximum of 100 RU at a flow rate of 10 μL/min using Biacore T200 (GE Healthcare, Chicago, IL). Serial dilutions of RTA in running buffer were injected at a rate of 50 μL/min. After each injection, the chip surface was regenerated with 10 mM glycine-HCl pH 2.0 at 30 μL/min for 40-45 s. Sensorgrams were normalized by subtracting baseline RU values from a reference flow cell (absent captured V_H_H) and analyzed by fitting the data to the 1:1 Langmuir binding model using the Biacore T200 Evaluation Software (GE Healthcare). Plots were created by importing the sensorgram data into GraphPad Prism 8.2 software.

### Vero cell culture, transfection, and lysis

Vero cells (ATCC, Manassas, VA) were cultured in Dulbecco’s minimal essential medium (DMEM) with fetal bovine serum (10% v/v) and penicillin/streptomycin at 37°C (5% CO_2_). Cells were transfected in 6-well (ELISAs) or 96-well plates (cytotoxicity assays) as recommended for Lipofectamine™ 3000 transfection protocol (Life Technologies, Carlsbad, CA). For ELISAs, cells were washed 48 h post-transfection twice in ice-cold PBS. RIPA Lysis Buffer (50 mM Tris-HCl pH7.5, 150 mM NaCl, 1% NP-40, 0.1% SDS, and 1% Na-deoxycholate; 150 μL) was added and cells were incubated for 5 minutes on ice with occasional gentle rocking to ensure well coverage. Cells were transferred to chilled 2 mL screw cap tubes with 1 mm glass beads by scraping and pipetting followed by homogenization at 5 m/s for 5 s. Cell lysates were replaced on ice for 30 min with one more homogenization step, followed by centrifugation at 15,700 x *g* for 10 min at 4°C.

For cytotoxicity assays, cells were transfected for 24 h before ricin treatment. Medium was aspirated from cells, replaced with 100 μL ricin toxin serially diluted in DMEM, and incubated at 37°C for 2 h. Ricin was then aspirated from cells and replaced with 100 μL DMEM. Cells were incubated for 48 h at 37°C. Viability was determined using Cell Titer-Glo® (Promega) and a Spectramax L Microplate Reader (Molecular Devices) and measured as a percentage of live control cells (transfected, but not treated with ricin).

### ELISAs

To detect intrabody levels, 96-well ELISA plates (Immulon™ 4HBX; ThermoFisher) were coated overnight at with 1 μg/mL SyH7 (anti-RTA mAb) (*50*) or 4 μg/mL asialofetuin (ASF; Sigma-Aldrich) in PBS. Plates were washed and blocked for 2 h at room temperature. Following block, ricin or RTA (1 μg/mL in PBS) was applied to ELISA plates for 1 h. Transfected cell lysate was added in triplicate and serially diluted across plate for 1 h. Plates were washed and HRP-conjugated anti-E-tag antibody (prepared following manufacturer’s guidelines; Bethyl Laboratories #A190-132P, Montgomery, TX) was applied for 1 h. Plates were washed and 100 μL TMB (SureBlue 3,3’,5,5’-tetramethylbenzidine; Kirkegaard & Perry Labs, Gaithersburg, MD) was added for 8-10 min followed by stop solution (1M phosphoric acid). The ELISA plates were analyzed using a SpectroMax250 spectrophotometer equipped with Softmax Pro 5.4.5 software (Molecular Devices, Sunnyvale, CA). Purified V_H_H protein with a C-terminal E-tag was used as a positive binding control for each transfection. Plates were washed following each step in PBS-Tween (0.1%). Cell lysates and secondary antibody were diluted in block buffer (wash buffer with 2% goat serum).

### Statistical analysis

Statistical analyses were performed using GraphPad Prism 8.2 software for Windows (San Diego, California). For *in vitro* translation assays, a Student’s t-test was used to compare to RTA control values. For intrabody cytotoxicity results, a two-way ANOVA was used with the Sidak post hoc test to compare transfected and LNP-vehicle control cell viabilities at each ricin concentration.

### Data Availability

All results associated with this study are included in the manuscript and supplementary information. As noted above, the crystal structures generated in this study were deposited in the PDB.

## Acknowledgements

We are grateful to Leslie Eisele and Renjie Song of the Wadsworth Center’s Biochemistry and Immunology Core facility for assistance with SPR. We thank the staff in the Media and Tissue Culture core for providing reagents, the Applied Genomics Technologies core for DNA sequencing and the Advance Light Microscopy core for assistance with confocal microscopy. We thank Dylan Ehrbar (Wadsworth Center) for assistance with statistical analysis, members of the Mantis lab for technical assistance, and Beth Cavosie for administrative support. We thank Dr. Chuck Shoemaker (Tufts University) and Dr. Anne Messer (Neural Stem Cell Institute) for guidance and advice with the intrabody studies. Finally, we gratefully acknowledge the 24-ID-C and 24-ID-E beamline staff at the Advanced Photon Source and beamline 8.2.2 at the Advanced Light Source, Berkeley Lab for their assistance in data collection.

## Funding

This work was supported by Contract No. HHSN272201400021C from the National Institutes of Allergy and Infectious Diseases. The content is solely the responsibility of the authors and does not necessarily represent the official views of the National Institutes of Health. The funders had no role in study design, data collection and analysis, decision to publish, or preparation of the manuscript.

## Author Contributions

MJR, solved, analyzed and annotated V_H_H-RTA crystal structures; SAD and CMTN assisted with structural biology; TC performed SPR, IVT and intrabody studies; XL and NT performed and analyzed depurination studies; DJV identified the V_H_Hs described in this manuscript, performed toxin neutralization assays and the V_H_H alignment analysis; MJR, TC, DJV and NJM wrote and edited the manuscript; NJM was responsible for project administration and funding acquisition.

## Competing interests

The authors have no financial or other competing interests to declare.

## Data and materials availability

The DNA sequences encoding the V_H_Hs described within this manuscript will be deposited into GenBank.

## References

1. M. de Virgilio, A. Lombardi, R. Caliandro, M. S. Fabbrini, Ribosome-inactivating proteins: from plant defense to tumor attack. Toxins (Basel) 2, 2699–2737 (2010).

2. T. Berger et al., Toxins as biological weapons for terror-characteristics, challenges and medical countermeasures: a mini-review. Disaster Mil Med 2, 7 (2016).

3. P. Grela, M. Szajwaj, P. Horbowicz-Drozdzal, M. Tchorzewski, How Ricin Damages the Ribosome. Toxins (Basel) 11, (2019).

4. N. Sowa-Rogozinska, H. Sominka, J. Nowakowska-Golacka, K. Sandvig, M. Slominska-Wojewodzka, Intracellular Transport and Cytotoxicity of the Protein Toxin Ricin. Toxins (Basel) 11, (2019).

5. A. Sapoznikov et al., Early disruption of the alveolar-capillary barrier in a ricin-induced ARDS mouse model: neutrophil-dependent and -independent impairment of junction proteins. Am J Physiol Lung Cell Mol Physiol, (2018).

6. Y. Endo, K. Tsurugi, RNA N-glycosidase activity of ricin A-chain. Mechanism of action of the toxic lectin ricin on eukaryotic ribosomes. J Biol Chem 262, 8128–8130 (1987).

7. W. Montfort et al., The three-dimensional structure of ricin at 2.8 A. Journal of Biological Chemistry. 262, 5398–5403 (1987).

8. M. P. Ready, Y. Kim, J. D. Robertus, Site-directed mutagenesis of ricin A-chain and implications for the mechanism of action. Proteins 10, 270–278 (1991).

9. D. J. Miller et al., Structure-based design and characterization of novel platforms for ricin and shiga toxin inhibition. J. Med. Chem. 45, 90–98 (2002).

10. Y. Endo, K. Tsurugi, The RNA N-glycosidase activity of ricin A-chain. The characteristics of the enzymatic activity of ricin A-chain with ribosomes and with rRNA. J Biol Chem 263, 8735–8739 (1988).

11. Y. P. Pang et al., Small-molecule inhibitor leads of ribosome-inactivating proteins developed using the doorstop approach. PLoS One 6, e17883 (2011).

12. K. Jasheway, J. Pruet, E. V. Anslyn, J. D. Robertus, Structure-based design of ricin inhibitors. Toxins (Basel) 3, 1233–1248 (2011).

13. B. Stechmann et al., Inhibition of retrograde transport protects mice from lethal ricin challenge. Cell 141, 231–242 (2010).

14. P. G. Wahome, S. Ahlawat, N. J. Mantis, Identification of small molecules that suppress ricin-induced stress-activated signaling pathways. PLoS One 7, e49075 (2012).

15. A. Desmyter, S. Spinelli, A. Roussel, C. Cambillau, Camelid nanobodies: killing two birds with one stone. Curr Opin Struct Biol 32, 1–8 (2015).

16. J. Dong et al., A single-domain llama antibody potently inhibits the enzymatic activity of botulinum neurotoxin by binding to the non-catalytic alpha-exosite binding region. J Mol Biol 397, 1106–1118 (2010).

17. J. M. Tremblay et al., Camelid single domain antibodies (VHHs) as neuronal cell intrabody binding agents and inhibitors of Clostridium botulinum neurotoxin (BoNT) proteases. Toxicon 56, 990–998 (2010).

18. C. E. Vrentas et al., A Diverse Set of Single-domain Antibodies (VHHs) against the Anthrax Toxin Lethal and Edema Factors Provides a Basis for Construction of a Bispecific Agent That Protects against Anthrax Infection. J Biol Chem 291, 21596–21606 (2016).

19. E. De Genst et al., Molecular basis for the preferential cleft recognition by dromedary heavy-chain antibodies. Proc Natl Acad Sci U S A 103, 4586–4591 (2006).

20. M. Lauwereys et al., Potent enzyme inhibitors derived from dromedary heavy-chain antibodies. EMBO J 17, 3512–3520 (1998).

21. S. K. Angalakurthi et al., A Collection of Single-Domain Antibodies that Crowd Ricin Toxin’s Active Site. Antibodies (Basel) 7, (2018).

22. D. J. Vance et al., High-Resolution Epitope Positioning of a Large Collection of Neutralizing and Nonneutralizing Single-Domain Antibodies on the Enzymatic and Binding Subunits of Ricin Toxin. Clin Vaccine Immunol 24, (2017).

23. Y. Zhou, X. P. Li, J. N. Kahn, N. E. Tumer, Functional Assays for Measuring the Catalytic Activity of Ribosome Inactivating Proteins. Toxins (Basel) 10, (2018).

24. A. L. Marschall, S. Dubel, T. Boldicke, Specific in vivo knockdown of protein function by intrabodies. MAbs 7, 1010–1035 (2015).

25. M. E. Fraser, M. M. Chernaia, Y. V. Kozlov, M. N. James, Crystal structure of the holotoxin from Shigella dysenteriae at 2.5 A resolution. Nat Struct Biol 1, 59–64 (1994).

26. C. Hamers-Casterman et al., Naturally occurring antibodies devoid of light chains. Nature 363, 446–448 (1993).

27. P. G. Wahome, J. D. Robertus, N. J. Mantis, Small-molecule inhibitors of ricin and Shiga toxins. Curr Top Microbiol Immunol 357, 179–207 (2012).

28. I. Achour et al., Tetrameric and homodimeric camelid IgGs originate from the same IgH locus. J Immunol 181, 2001–2009 (2008).

29. J. M. Pruet et al., 7-Substituted pterins provide a new direction for ricin A chain inhibitors. Eur J Med Chem 46, 3608–3615. (2011).

30. J. G. Park, J. N. Kahn, N. E. Tumer, Y. P. Pang, Chemical Structure of Retro-2, a Compound That Protects Cells against Ribosome-Inactivating Proteins. Sci Rep 2, 631 (2012).

31. A. Desmyter et al., Crystal structure of a camel single-domain VH antibody fragment in complex with lysozyme. Nat Struct Biol 3, 803–811 (1996).

32. A. Desmyter et al., Three camelid VHH domains in complex with porcine pancreatic alpha-amylase. Inhibition and versatility of binding topology. J Biol Chem 277, 23645–23650 (2002).

33. S. Muyldermans, Nanobodies: natural single-domain antibodies. Annu Rev Biochem 82, 775–797 (2013).

34. P. Acharya et al., Heavy chain-only IgG2b llama antibody effects near-pan HIV-1 neutralization by recognizing a CD4-induced epitope that includes elements of coreceptor- and CD4-binding sites. J Virol 87, 10173–10181 (2013).

35. M. Strauss, L. Schotte, B. Thys, D. J. Filman, J. M. Hogle, Five of Five VHHs Neutralizing Poliovirus Bind the Receptor-Binding Site. J Virol 90, 3496–3505 (2016).

36. T. R. Transue, E. De Genst, M. A. Ghahroudi, L. Wyns, S. Muyldermans, Camel single-domain antibody inhibits enzyme by mimicking carbohydrate substrate. Proteins 32, 515–522 (1998).

37. T. Boldicke, Single domain antibodies for the knockdown of cytosolic and nuclear proteins. Protein Sci 26, 925–945 (2017).

38. R. H. Argent et al., Ribosome-mediated folding of partially unfolded ricin A-chain. J Biol Chem 275, 9263–9269 (2000).

39. M. Thran et al., mRNA mediates passive vaccination against infectious agents, toxins, and tumors. EMBO Mol Med 9, 1434–1447 (2017).

40. M. J. Rudolph et al., Crystal structures of ricin toxin’s enzymatic subunit (RTA) in complex with neutralizing and non-neutralizing single-chain antibodies. J Mol Biol 426, 3057–3068 (2014).

41. P. G. Wahome, N. J. Mantis, High-throughput, cell-based screens to identify small-molecule inhibitors of ricin toxin and related category b ribosome inactivating proteins (RIPs). Curr Protoc Toxicol Chapter 2, Unit 2.23 (2013).

42. X. P. Li, P. C. Kahn, J. N. Kahn, P. Grela, N. E. Tumer, Arginine residues on the opposite side of the active site stimulate the catalysis of ribosome depurination by ricin A chain by interacting with the P-protein stalk. J Biol Chem 288, 30270–30284 (2013).

43. M. Pierce, J. N. Kahn, J. Chiou, N. E. Tumer, Development of a quantitative RT-PCR assay to examine the kinetics of ribosome depurination by ribosome inactivating proteins using Saccharomyces cerevisiae as a model. RNA 17, 201–210 (2011).

44. Z. Otwinowski, W. Minor, Processing of x-ray diffraction data collected in oscillation mode. . Methods in Enzymology 276, 307–326 (1997).

45. M. D. Winn et al., Overview of the CCP4 suite and current developments. Acta Crystallogr D Biol Crystallogr 67, 235–242 (2011).

46. A. J. McCoy et al., Phaser crystallographic software. J Appl Crystallogr 40, 658–674 (2007).

47. R. J. Morris, A. Perrakis, V. S. Lamzin, ARP/wARP and automatic interpretation of protein electron density maps. . Methods Enzymol 374, 229–244 (2003).

48. P. Emsley, B. Lohkamp, W. G. Scott, K. Cowtan, Features and development of Coot. Acta Crystallogr D Biol Crystallogr 66, 486–501 (2010).

49. P. D. Adams et al., PHENIX: a comprehensive Python-based system for macromolecular structure solution. Acta Crystallogr D Biol Crystallogr 66, 213–221 (2010).

50. R. T. t. Toth et al., High-Definition Mapping of Four Spatially Distinct Neutralizing Epitope Clusters on RiVax, a Candidate Ricin Toxin Subunit Vaccine. Clin Vaccine Immunol 24, (2017).

