## Supplemental Information for "Intracellular Neutralization of Ricin Toxin by Single Domain Antibodies Targeting the Active Site Pocket"

### Supplemental Tables

**Table S1. $V_H$H Binding Kinetics for RTA and Correlation to *in vitro* Neutralization**

| $V_H$H | $K_{on}$ (1/Ms) | $K_{off}$ (1/s) | $K_D$ (nM)[a] |
|---|---|---|---|
| V2A11 | $2.2 \times 10^{6}$ | $6.8 \times 10^{-4}$ | 0.31 ± 0.01 |
| V6H8 | $5.3 \times 10^{6}$ | $2.1 \times 10^{-2}$ | 4.0 ± 1.9 |
| V8E6 | $1.6 \times 10^{6}$ | $6.9 \times 10^{-4}$ | 0.42 ± 0.12 |
| V6A7 | $1.7 \times 10^{6}$ | $7.1 \times 10^{-3}$ | 4.2 ± 0.72 |
| V6A6 | $3.7 \times 10^{5}$ | $4.0 \times 10^{-3}$ | 10.9 ± 5.3 |
| V2G10 | $2.2 \times 10^{6}$ | $6.2 \times 10^{-4}$ | 0.28 ± 0.24 |
| V6D4 | $1.4 \times 10^{7}$ | $1.4 \times 10^{-1}$ | 9.5 ± 2.3 |

[a], Binding affinities for RTA were determined by SPR, as described in the Materials and Methods. Values represent the mean of three replicates with standard deviations.

**Table S2. Data collection associated with RTA-V$_H$H X-ray crystal structures**

| RTA Complex: | V2A11 | V6H8 | V8E6 | V6A7 | V6A6 | V2G10 | V6D4 |
|---|---|---|---|---|---|---|---|
| APS Beamline | ALS-501 | 24-ID-E | 24-ID-C | 24-ID-E | 24-ID-C | 24-ID-E | 24-ID-C |
| $d_{min}$ (Å) | 1.8 | 1.7 | 2.0 | 2.5 | 1.9 | 2.1 | 2.1 |
| wavelength (Å) | 0.977 | 0.979 | 0.979 | 0.979 | 0.979 | 0.979 | 0.979 |
| No. reflections | 227994 | 1853464 | 1502572 | 521771 | 5587958 | 771102 | 617202 |
| Ave. redundancy[a] | 1.1(1.1) | 10.9(6.9) | 1.7(1.6) | 7.2(3.9) | 2.8(2.0) | 3.0(2.3) | 1.9(1.7) |
| (*I*)/(δ)[a] | 26.2(2.1) | 33.6(1.7) | 16.1(1.1) | 29.1(0.8) | 8.4(1.1) | 17.4(0.72) | 10.6(0.9) |
| Completeness[a] (%) | 96.8(96.4) | 100(100) | 96.2(91.8) | 99.7(94.8) | 99.9(99.4) | 99.2(90.0) | 99.7(97.7) |
| $R_{merge}$[a,b] (%) | 3.8(42.1) | 11.5(256.3) | 7.1(57.7) | 6.1(122.5) | 25.5(233.7) | 7.2(128.6) | 12.4(104.9) |
| CC*[c] | (0.89) | (0.90) | (0.90) | (0.90) | (0.80) | (0.81) | (0.83) |
| **Refinement** | | | | | | | |
| Bragg spacings (Å) | 43.2-1.8 | 44.1-1.7 | 45.7-2.0 | 46.6-2.5 | 46.4-2.1 | 48.1-2.1 | 46.4-2.1 |
| Space group | P1 | $P6_522$ | $P2_1$ | I422 | C2 | C2 | $P2_1$ |
| Cell parameters: *a,b,c* (Å) / α,β,γ (°) | 38.7, 47.2, 58.4, / 100.3, 97.3, 109.7 | 90.3, 90.3, 204.0 | 59.7, 77.9, 92.6 / b = 98.9 | 102.6, 102.6, 156.8 | 123.4, 112.9, 93.1 / b=127.5 | 193.9 50.3 111.7 / b=124.7 | 57.9, 78.6, 103.9 / b = 94.5 |
| $R^d$ / $R_{free}^e$ (%) | 18.7/24.7 | 17.1/20.8 | 18.9/21.8 | 21.1/26.0 | 20.7/24.4 | 21.3/24.7 | 18.9/22.8 |
| No. of reflections | 35816 | 52032 | 54868 | 14930 | 79403 | 50791 | 53533 |
| No. of waters | 272 | 433 | 324 | 14 | 833 | 201 | 301 |
| Rmsd bond length (Å) | 0.006 | 0.011 | 0.006 | 0.003 | 0.010 | 0.002 | 0.003 |
| Rmsd bond angle (°) | 0.97 | 1.264 | 0.797 | 0.732 | 1.057 | 0.622 | 0.671 |
| Ramachandran favored/allowed[f] (%) | 99.5 / 100 | 98.4 / 100 | 98.4 / 100 | 96.5 / 99.7 | 98.4 / 99.9 | 97.4 / 100 | 98.4 / 100 |
| **PDB code** | **6OBC** | **6OBE** | **6OBG** | **6OBM** | **6OBO** | **6OCA** | **6OCD** |

[a] Values in outermost shell are given in parentheses.

[b] $R_{merge} = (\sum | I_i - <I_i> |)/\sum | I_i |$, where $I_i$ is the integrated intensity of a given reflection.

[c] CC*=□2CC1/2/1+CC1/2 , where CC1/2 is the correlation coefficient of two split data sets each derived by averaging half of the observations for a given reflection.

[d] $R = \sum | |F_o| - |F_c| | / \sum | |F_o| |$, where $F_o$ and $F_c$ denote observe and calculated structure factors, respectively.

[e] $R_{free}$ was calculated using 5% of data excluded from refinement.

[f] Calculated using Molprobity.

**Table S3. Categorization of CDR3 Conformations**

| | $V_HH$ | PDB ID |
|---|---|---|
| Extended | JIY-D10 | 4LGR |
| | JIY-G11 | 4LHJ |
| | V2A11 | 6OBC |
| | V6A6 | 6OBO |
| | V6A7 | 6OBM |
| | V6H8 | 6OBE |
| | V8E6 | 6OBG |
| Pinned | JIV-G12 | 4LGS |
| | JIY-A7 | 4LHQ |
| | JIY-E5 | 4LGP |
| | JIV-F5 | 4Z9K |
| | JNM-F8 | 5E1H |
| | JIY-E1 | 5BOZ |
| | V1C7 | 5J56 |
| | V5E1 | 5J57 |
| | JPF-A9 | 6CWG |
| | V2G10 | 6OCA |
| | V6D4 | 6OCD |

#### Supplemental Figure Legends

**Figure S1. Epitope localization and toxin-neutralizing activities of seven $V_HH$s in this report.** (**Panel A**) The epitopes recognized by the seven RTA-specific $V_HH$s described in this report were positioned within subcluster 3.1 (left) or subcluster 3.3 (right), as described Angalakurthi et al., 2018. Colored are RTA (light grey), RTB (charcoal) and the active site (yellow), plus RTA secondary structures α-helix C (red), α-helix G (green), and α-helix B (blue). (**Panel B**) In a Vero cell cytotoxicity assay (see Materials and Methods), only V6D4 (subcluster 3.3) afforded any detectable toxin-neutralizing activity with ~60% cell viability and estimated $IC_{50}$ of ~200 nM. (**Panel C**) The amino acid sequences of the seven $V_HH$s described in this report were aligned using Clustal Omega. CDR1 (blue), CDR2 (yellow) and CDR3 (red) are colored accordingly. Black asteriks below the sequence denote sequence identity, black colons and black periods indicate higher and lower levels of sequence similarity, respectively. Clustal Omega: https://www.ebi.ac.uk/Tools/msa/clustalo/

**Figure S2. Representative sensorgrams associated with $V_HH$ binding to RTA.** Rabbit polyclonal anti-E-tag antibodies were covalently linked to a series S CM5 sensor chip using a Biacore T200. Each of the seven $V_HH$s were captured via their C-terminal E-tags up to a maximum of 100 RU. RTA was injected at the indicated concentrations at a flow rate of 50 μL/min in running buffer. The chip surface was regenerated at pH 2.0 for 40-45 s following each RTA injection. Each experiment was repeated three times. Shown is a representative sensorgram for each $V_HH$. The data were fit using a 1:1 Langmuir binding model.

**Figure S3. ELISA-based detection of $V_HH$ intrabodies following transient transfection of Vero cells. (A)** Schematic of workflow. Vero cells were seeded in 6-well plates and then transiently transfected with $V_HH$-encoding pcDNA3.1 plasmids. Two days later, the cells were lysed. The lysates were serially diluted and applied to RTA- or ricin-captured ELISA plates as indicated. $V_HH$s, with C-terminal E-tags, were detected with HRP-conjugated anti-E-tag antibody. **(B)** SyH7 (anti-RTA mAb) was coated on the ELISA plates prior to RTA capture. **(D)** ASF was coated on the ELISA plates prior to ricin capture. **(F)** SyH7 was coated on ELISA plates prior to ricin capture. **(C, E, F)** Control experiments for panels B, D, and F, respectively, in which purified $V_HH$s with C-terminal E-tags serially diluted in block buffer were applied to RTA- or ricin-captured ELISA plates and detected with HRP-conjugated anti-E-tag antibody. Each panel (**B-G**) represents a single experiment done in triplicate.

**Figure S4. RTA secondary elements.** Ribbon diagram of RTA in green (PDB ID **1RTC**) with all secondary structures labeled.

**Figure S5. Illustration of the different $V_HH$ binding modes with RTA's active site.** Superpositioned Cα-traces of: (**Panel A**) Close-up of the superposed CDR3 regions of V2A11 (cyan) and V6A7 (gray) depicting their different configurations. Side chains are drawn as sticks and color coordinated to the main chain color; (**Panel B**) Superpositioned Cα-traces of RTA-V2A11 (green-cyan) with RTA-V6A6 (gray-gray). (**Panel C**) RTA-V2A11 (green-cyan) with RTA-V6H8 (gray-gray); (**Panel D**) RTA-V6A6 (green-cyan) with RTA-V6A7 (gray-gray); (**Panel E**) RTA-V6A6 (green-cyan) and RTA-V8E6 (gray-gray).

**Figure S6. VHH Mode 1A and Mode 1B interactions with the RTA active site residues.** (Panel A) Close-up of the superposed active sites of RTA (green) bound to V2A11 (red) and RTA (gray) bound to the substrate analog adenosine monophosphate depicting the different configurations of key RTA active site residue Tyr-80. Side chains are drawn as sticks and color coordinated to the main chain color. Adenosine monophosphate is also drawn as sticks and colored by atom type with oxygen red, nitrogen blue, carbon gray, and sulfur orange (**PDB 3HIO**). (B) Close-up of the superposed active site of RTA (green) bound to V2A11 (red) and RTA (gray) bound to the tetranucleotide substrate analog (carbons gray and nitrogens blue) depicting the different distances of RTA catalytic residue Arg-180 from the N-3 atom within the adenine ring of the tetranucleotide. V2A11 CDR3 Ser-103 and Arg-180 in RTA are drawn as sticks and color coordinated to their respective main chain colors. The hydrogen bond between CDR3 Ser-103 and RTA Arg-180 is represented as red dashes as are the different distances between Arg-180 to the N-3 atom within the adenine ring of the tetranucleotide.

**Figure S7**. **Influence of CDR1 and CDR2 configuration on binding affinity.** (**Panel A**) Superpositioned Cα-traces of RTA-V2A11 complex with RTA-V6H8. The CDR1 element in V2A11 is colored blue while CDR1 region of V6H8 is colored dark gray. (**Panel B**) Superpositioned Cα-traces of V6A6 with V6A7, V8E6, and V6H8 all bound to RTA showing their similar configuration. In the RTA-V6H8 complex, RTA is colored green and V6H8 is colored cyan with its CDR1 element colored blue. All other RTA-VHH complexes are colored gray. (**Panel C**) The CDR2 segment in V2A11 is colored yellow while CDR2 element of V6H8 is colored dark gray. RTA is colored green, the framework residues in V2A11 are colored in cyan, and light gray in V6H8 colored light gray in panels A and B. Important residues involving paratope-epitope interactions are drawn as sticks and color coordinated to their respective main chain color. Hydrogen bonds and salt bridges are represented as red dashes.

**Figure S8**. **Additional $V_H$H binding modes with RTA's active site.** The Cα-traces of RTA (green) in complex with (**Panel A**) V2A11, V2G10, and V6D4. $V_H$Hs are colored in cyan; CDRs 1, 2, and 3 are colored blue, yellow, and red, respectively. Arrows illustrate the direction of the ~99° rotation of V2G10 and the ~165° rotation of V6D4, each relative to V2A11. (**Panel B**) Superpositioned Cα-traces of V2A11 onto V2G10 and V6D4 demonstrating V2A11's CDR3 conformation relative to the other two $V_H$Hs. CDR3 elements are colored red. V2G10 residues involved in hydrophobic interactions between CDR3 and FR are drawn as sticks and color coordinated to their respective main chain color. The disulfide bond between residues Cys50 and Cys105 of V6D4 is shown in stick representation and colored magenta.

**Figure S9**. **Different interactions of V6D4, V2A11 and V6A6 with RTA's α-helix B.** Four different $V_H$H (cyan) interactions with RTA's α-helix B, a known neutralizing hotspot on ricin toxin. (**Panel A**) V6D4 interactions with several solvent-exposed residues of α-helix B.(**Panels B-C**) V2A11 and V6A6 CDR3 interactions with several less solvent exposed residues within α-helix B; (D) Binding of the potent neutralizing $V_H$H E5 with the solvent accessible region of RTA's α-helix B. The hydrogen bond between E5 CDR3 residue Arg-104 and RTA Thr-105 is represented as red dashes. RTA (green) and the $V_H$Hs (cyan) are drawn as ribbon diagrams. CDRs 1, 2, and 3 are colored blue, yellow, and red, respectively. Key side chains are drawn as sticks and color coordinated to the main chain color. Hydrogen bonds are represented as red dashes.

**Figure S10. Extended and pinned configurations of alpaca $V_HH$ CDR3s elements.** Superpositioned Cα-traces of 18 alpaca-derived (*Vicuga pacos*) $V_HH$ crystal structures listed in **Table S3**. Within this sampling, the CDR3 elements assume two distinct conformations: extended and pinned. (**Panel A**) In the extended conformations, the CDR3 element (red) projects upwards from β-strand F and away from CDRs 1 (blue) and 2 (yellow), before returning downward to β-strand G. (**Panel B**) In the pinned or "reefed" conformation, the CDR3 elements (red) fold back onto CDRs 1 (blue) and 2 (yellow), ultimately making contact with vestigial FR2 residues formerly involved in VL interactions, before connecting with β-strand G. Cysteine residues involved in non-canonical disulfides are colored magenta.

### Figure S1

**A**

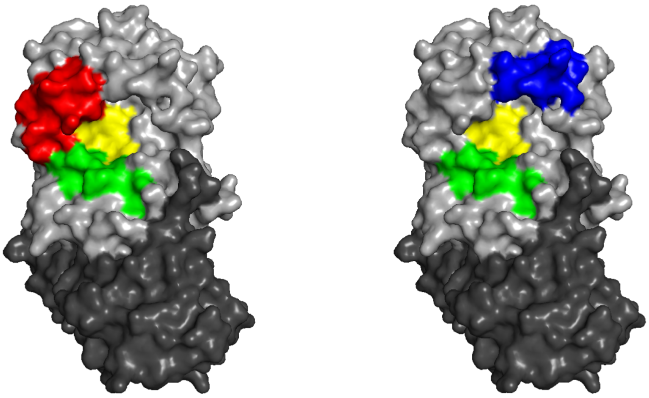


**B**

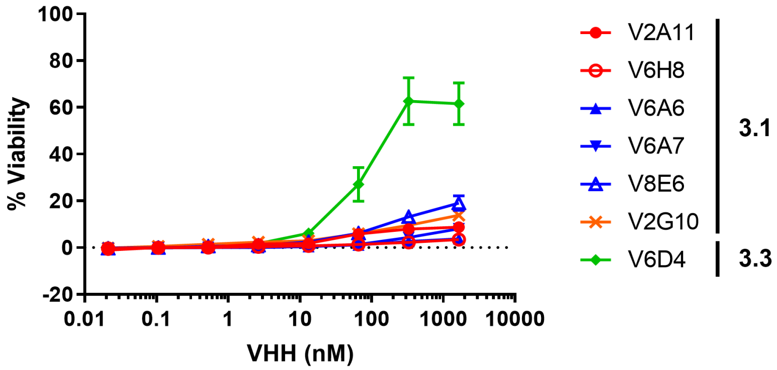


**C**

```
V2G10     QVQLVET-GGGVVQAGGSLRLSCVASGRTFSVSGRTFSDHGLGWFRQAPGKEREFVGSIS 59
V6D4      QLQLVET-GGGLVQSGGSLRLSCAASGFTL-------DNYNIGWFRQAPGKEYGGVSCIS 52
V2A11     QVQLAET-GGGLVQPGGARTLSCAASESIS-------SFYFMGWYRQAPGKPRELVAEIS 52
V6H8      QLQLVET-GGGLVQAGGSLRLSCAASGSIF-------SMHAMGWFRQAPGRERELVAVAP 52
V6A7      QVQLVETGGGGLVQAGGSLRLSCAASGSIS-------SLNAMGWYRQAPGKERELVADIS 53
V6A6      QVQLAET-GGGLAQAGGSLRLSCAASGSIF-------SINAMGWYRQAPGKERELVADIS 52
V8E6      QVQLVET-GGGLVQPGGSLRLSCAASGSIF-------SINAMGWYRQAPGKERELVADIS 52
          *:**.** ***:.* **:   ***.**          .  :**:*****:    *.

V2G10     WSVDGDATYYTDLANSVKGRFTISGVNAKNTVYLQMNSLKPEDTAVYYCAAGLRGGTYAR 119
V6D4      SSD-----GSTYYADSVKGRFTISRDNAKNTVYLQMNNLKPEDTDVYYCAATKY--GSSC 105
V2A11     -NY-----GRTDYGDSLKGRFTISRDNAANTVNLQMNNLAPEDTALYYCNARKW--ER-- 102
V6H8      -TG-----RPSDYADFAKGRFTISRDNAKNTVSLQMHSLEPEDTAVYYCNAQLW--ER-- 102
V6A7      -AS-----GRTNYADSVKGRFTISRDNAKNTVSLQMNSLKPEDTAVYYCNAVGG--TY-- 103
V6A6      -GS-----GRTNYADSVKGRFTISRDNAKNTVSLQMNSLKPEDTAVYYCNVVGG--SY-- 102
V8E6      -SS-----GRINEADSVKGRFTISRDNAKNTVYLQMNSLKPEDTAVYYCNVLAG--SH-- 102
                       .:  *******  ** *** ***:.* **** :*** .

V2G10     TIYEYDYWGQGTQVTVSLEPKTPKPQ   145
V6D4      PIRPYDYWGQGTQVTVSSAHHSEDPS   131
V2A11     -SVLEDYWGQGTQVTVSSEPKTPKPQ   127
V6H8      -YVLNDYWGQGTQVTVSSEPKTPKPQ   127
V6A7      YYDEYDYWGQGTQVTVSSAHHSEDPS   129
V6A6      YYDEYNYWGQGTQVTVSSEPKTPKPQ   128
V8E6      YYDEYEYWGQGTQVTVSSEPKTPKPQ   128
               :***********   :: .*.
```

#### Figure S2

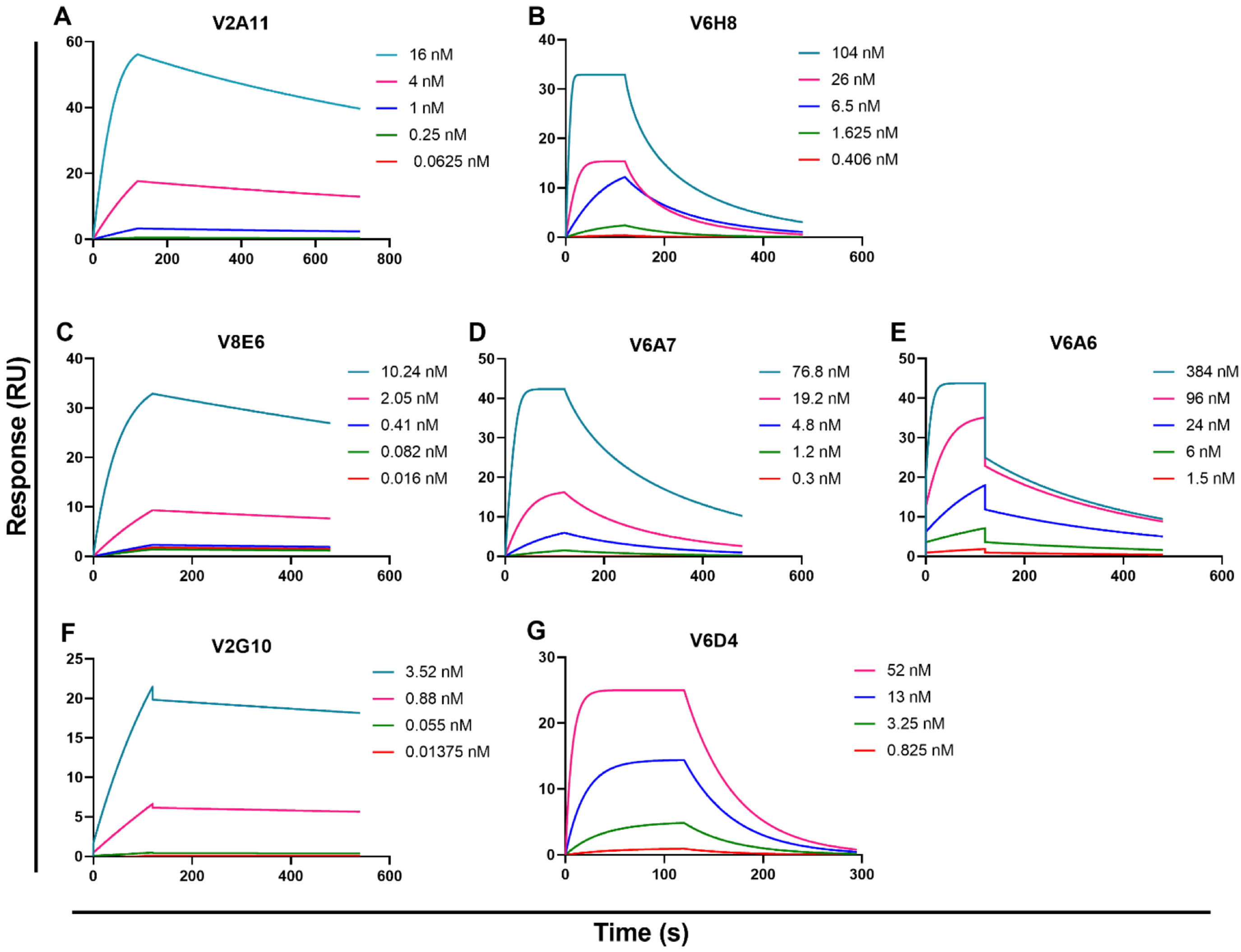
A
V2A11
16 nM
4 nM
1 nM
0.25 nM
0.0625 nM
B
V6H8
104 nM
26 nM
6.5 nM
1.625 nM
0.406 nM
C
V8E6
10.24 nM
2.05 nM
0.41 nM
0.082 nM
0.016 nM
D
V6A7
76.8 nM
19.2 nM
4.8 nM
1.2 nM
0.3 nM
E
V6A6
384 nM
96 nM
24 nM
6 nM
1.5 nM
F
V2G10
3.52 nM
0.88 nM
0.055 nM
0.01375 nM
G
V6D4
52 nM
13 nM
3.25 nM
0.825 nM
Response (RU)
Time (s)

**Figure S3**

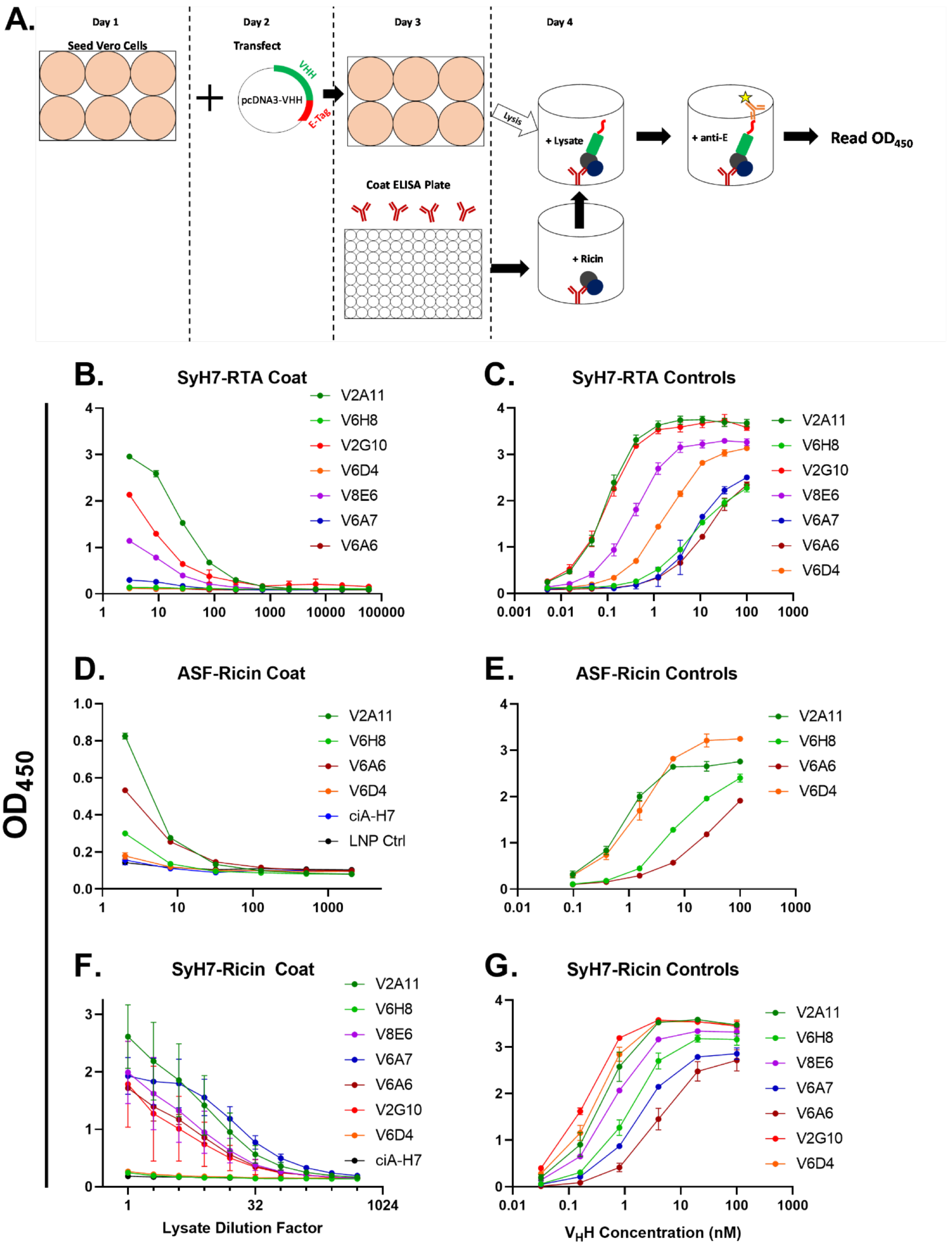
A.
Day 1
Seed Vero Cells
Day 2
Transfect
pcDNA3-VHH
VHH
E-Tag
Day 3
Coat ELISA Plate
Day 4
Lysis
+ Lysate
+ anti-E
Read OD450
+ Ricin
B.
SyH7-RTA Coat
V2A11
V6H8
V2G10
V6D4
V8E6
V6A7
V6A6
C.
SyH7-RTA Controls
V2A11
V6H8
V2G10
V8E6
V6A7
V6A6
V6D4
D.
ASF-Ricin Coat
V2A11
V6H8
V6A6
V6D4
ciA-H7
LNP Ctrl
E.
ASF-Ricin Controls
V2A11
V6H8
V6A6
V6D4
F.
SyH7-Ricin Coat
V2A11
V6H8
V8E6
V6A7
V6A6
V2G10
V6D4
ciA-H7
G.
SyH7-Ricin Controls
V2A11
V6H8
V8E6
V6A7
V6A6
V2G10
V6D4
OD450
Lysate Dilution Factor
VHH Concentration (nM)

**Figure S4**

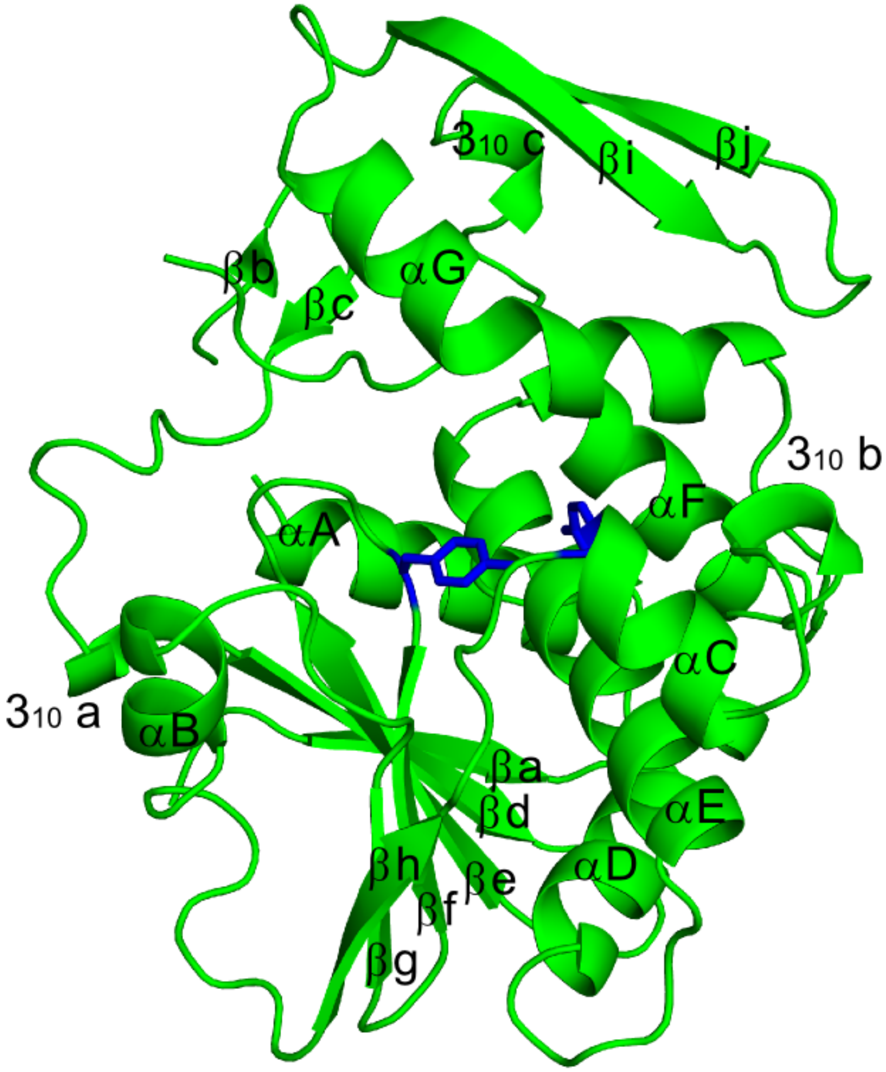

**Figure S5**

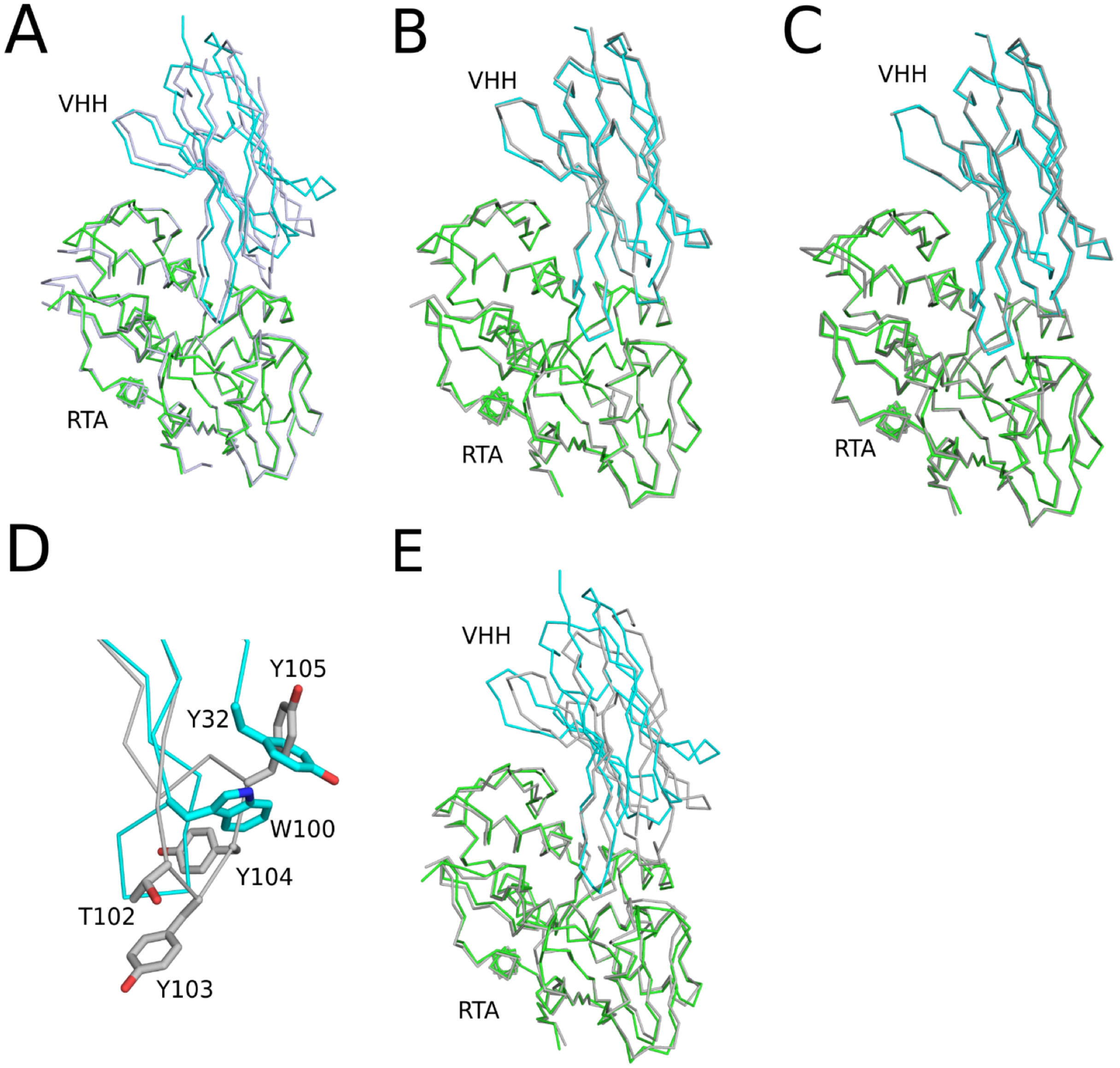

A
VHH
RTA
B
VHH
RTA
C
VHH
RTA
D
Y105
Y32
W100
Y104
T102
Y103
E
VHH
RTA

**Figure S6**

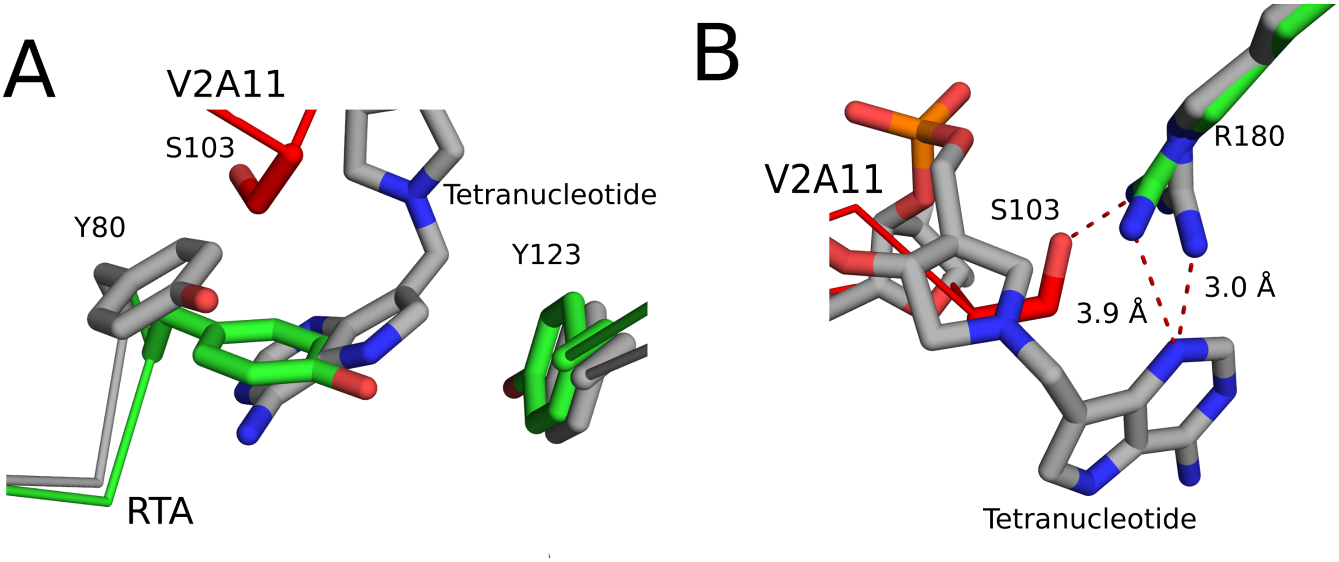
A
V2A11
S103
Tetranucleotide
Y80
Y123
RTA
B
R180
V2A11
S103
3.9 Å
3.0 Å
Tetranucleotide

**Figure S7**

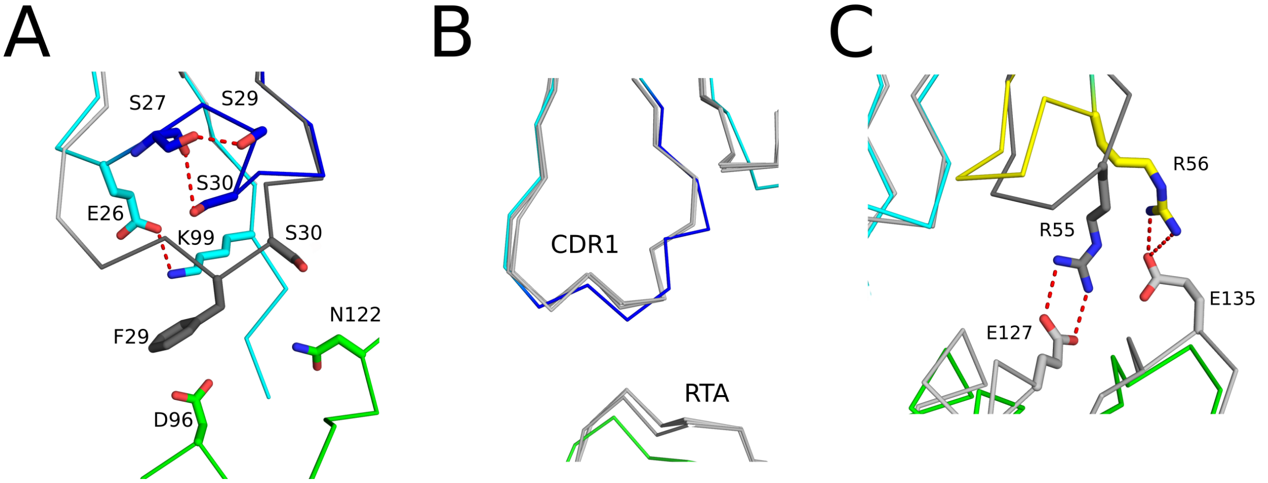
A
S27
S29
S30
E26
K99
S30
F29
N122
D96
B
CDR1
RTA
C
R56
R55
E135
E127

**Figure S8**

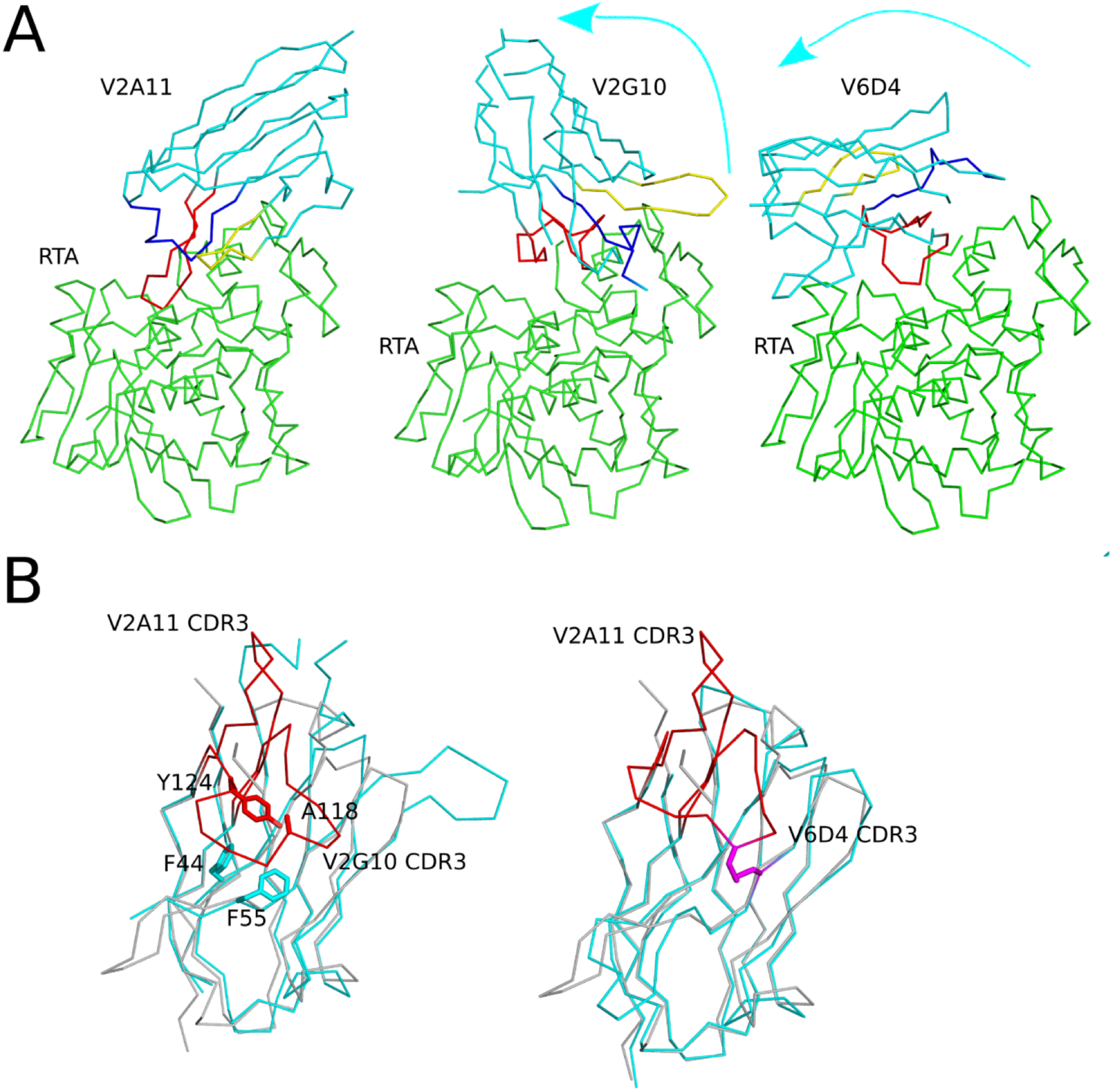
A
V2A11
V2G10
V6D4
RTA
RTA
RTA
B
V2A11 CDR3
Y124
A118
F44
V2G10 CDR3
F55
V2A11 CDR3
V6D4 CDR3

**Figure S9**

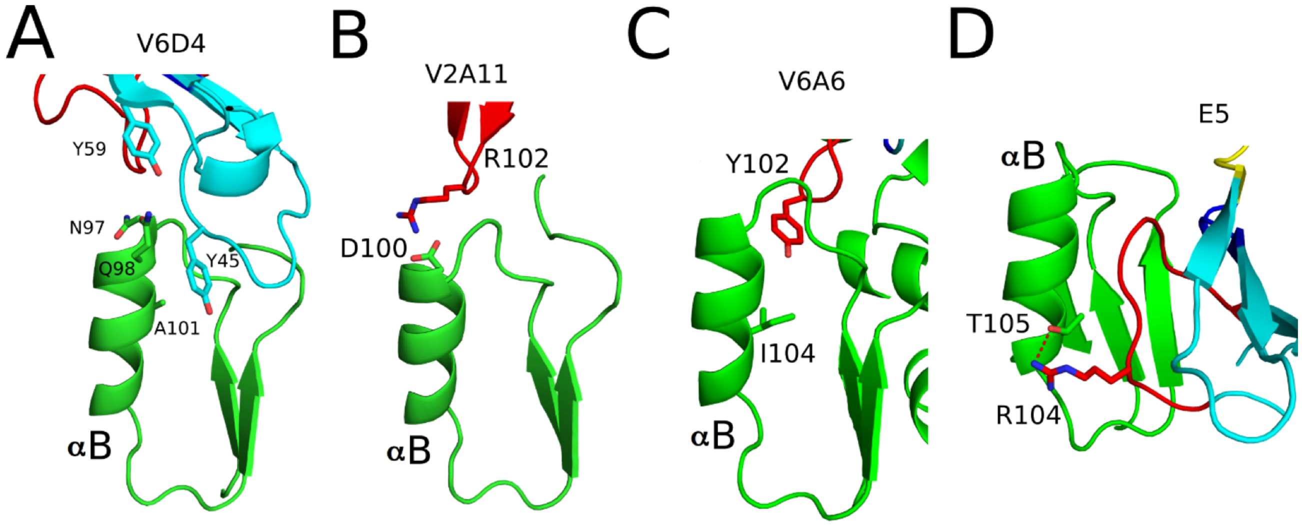
A
V6D4
Y59
N97
Q98
Y45
A101
αB
B
V2A11
R102
D100
αB
C
V6A6
Y102
I104
αB
D
E5
αB
T105
R104

**Figure S10**

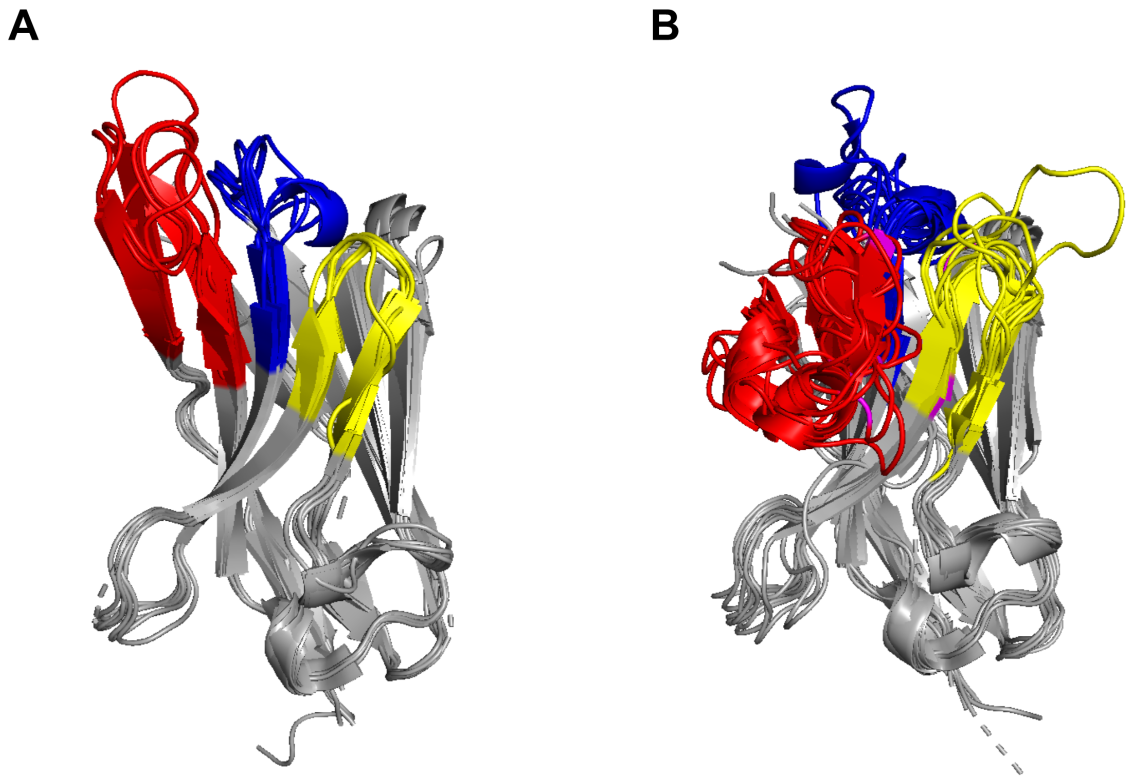
